# CTCF barrier breaking by ZFP661 promotes protocadherin diversity in mammalian brains

**DOI:** 10.1101/2023.05.08.539838

**Authors:** Jinpu Jin, Sherry Ralls, Elaine Wu, Gernot Wolf, Ming-An Sun, Danielle A. Springer, Rachel L. Cosby, Anna D. Senft, Todd S. Macfarlan

## Abstract

Mammalian brains are larger and more densely packed with neurons than reptiles, but the genetic mechanisms underlying the increased connection complexity amongst neurons are unclear. The expression diversity of clustered protocadherins (Pcdhs), which is controlled by CTCF and cohesin, is crucial for proper dendritic arborization and cortical connectivity in vertebrates. Here, we identify a highly-conserved and mammalian-restricted protein, ZFP661, that binds antagonistically at CTCF barriers at the *Pcdh* locus, preventing CTCF from trapping cohesin. ZFP661 balances the usage of Pcdh isoforms and increases Pcdh expression diversity. Loss of *Zfp661* causes cortical dendritic arborization defects and autism-like social deficits in mice. Our study reveals both a novel mechanism that regulates the trapping of cohesin by CTCF and a mammalian adaptation that promoted Pcdh expression diversity to accompany the expanded mammalian brain.

**One sentence summary:** ZFP661 blocks cohesin trapping by CTCF and increases protocadherin diversity for proper cortical dendritic arborization.

## Main Text

Mammals have enlarged brains relative to reptiles, with greater numbers of neurons and increased neuronal density, especially in the cerebrum and cerebellum (*1, 2*). The increased number and density of neurons in mammals likely necessitated evolutionary adaptations to allow neurites to properly identify each other and intermingle in a more crowded space during development. However, the genetic mechanisms that underlie the adaptations are not well understood. In vertebrates, the clustered protocadherin genes (Pcdhs), which encode cell adhesion molecules, play a central role as neuronal identity tags (*3-7*). Pcdhs have dozens of alternative isoforms, and only a selected subset of isoforms is expressed in individual neurons, producing a large diversity of isoform combinations (*5*). Neurites with the same/similar Pcdh combinations repel each other (*6-8*) whereas neurites with diverse Pcdhs can intermingle (*5, 6*). Repulsive signals amongst neurites are likely achieved from the formation of zipper-like chains mediated by isoform-specific homophilic interactions of Pcdh molecules, in such a way that isoform mismatch would terminate chain formation and reduce the chance of forming repulsive signals (*9-11*). At the transcriptional level, the combination of repression (*12, 13*) and stochastic demethylation (*14*) of alternative Pcdh promoters produces the diverse Pcdh isoforms that are accessible for activation in individual neurons. The enhancers downstream of the *Pcdh* locus further activate the expression of accessible Pcdh isoforms, dependent on CTCF and cohesin (*14-16*). CTCF and cohesin are two key players that control 3D genome structures and gene expression; ATP-dependent cohesin extrusion of chromatin promotes enhancer-promoter interactions, while CTCF acts as a barrier to stop (*i*.*e*., trap) cohesin extrusion and confine most enhancer-promoter interactions within the CTCF boundary (*17-21*). Whether the trapping of cohesin by CTCF can be regulated is also unclear.

Krüppel-associated box zinc finger proteins (KRAB-ZFPs) represent the largest family of transcription factors in mammals. Although many of these factors are species-restricted and bind to and repress lineage specific retrotransposons (*22-24*), a large number also emerged in the last common ancestor of mammals (*23, 25*). Some of these factors, like ZFP57/ZFP445 and ZFP568 play critical roles in mammalian-specific phenomenon like genomic imprinting and placental *Igf2* suppression (*26-28*), although the vast majority have gone unexplored. Many KRAB-ZFPs are preferentially expressed in human fetal brains (*29, 30*) and a small number have been shown to overlap CTCF and RAD21 (a subunit of cohesin complex) binding (*23*), indicating their potential roles in brain development and evolution by rewiring chromatin loops. Here, we identified a conserved KRAB-ZFP in Theria, ZFP661 (ZNF2 in humans), that binds to a small subset of CTCF sites genome-wide, including a cluster of CTCF sites at the *Pcdh* locus. Using mouse genetics, we uncovered a critical function for ZFP661 in modulating the trapping of cohesin at CTCF barriers, diversifying neuronal Pcdh expression and shaping mammalian brain development.

### ZFP661 is highly conserved in Theria and suppresses cohesin trapping at CTCF barriers without altering CTCF binding

To identify a candidate to explore the potential role of KRAB-ZFPs in rewiring 3D genome structures, we re-analyzed ChIP-seq data of 221 human KRAB-ZFPs (*23*) and found that 77.4% (171/221) of KRAB-ZFPs had binding peaks that significantly overlapped CTCF and RAD21 peaks (Fig. 1A). Amongst these KRAB-ZFPs, ZNF2 (known as ZFP661 in mice), was unique due to both the number and proportion of ChIP peaks that overlapped with CTCF and RAD21 (Fig. 1A). We determined that ZFP661 originated before the divergence of placental mammals and marsupials and that it is present in most (89/94) of the available therian genomes (Fig. 1B) according to a gain/loss tree at Ensembl. Notably both the DNA-binding domain (DBD) (90.1% identity across three representative species) and zinc fingerprint amino acids that are responsible for making specific DNA base contacts (97.2% identity) are highly conserved (fig. S1), suggesting it serves an important function in mammals.

**Fig. 1.**
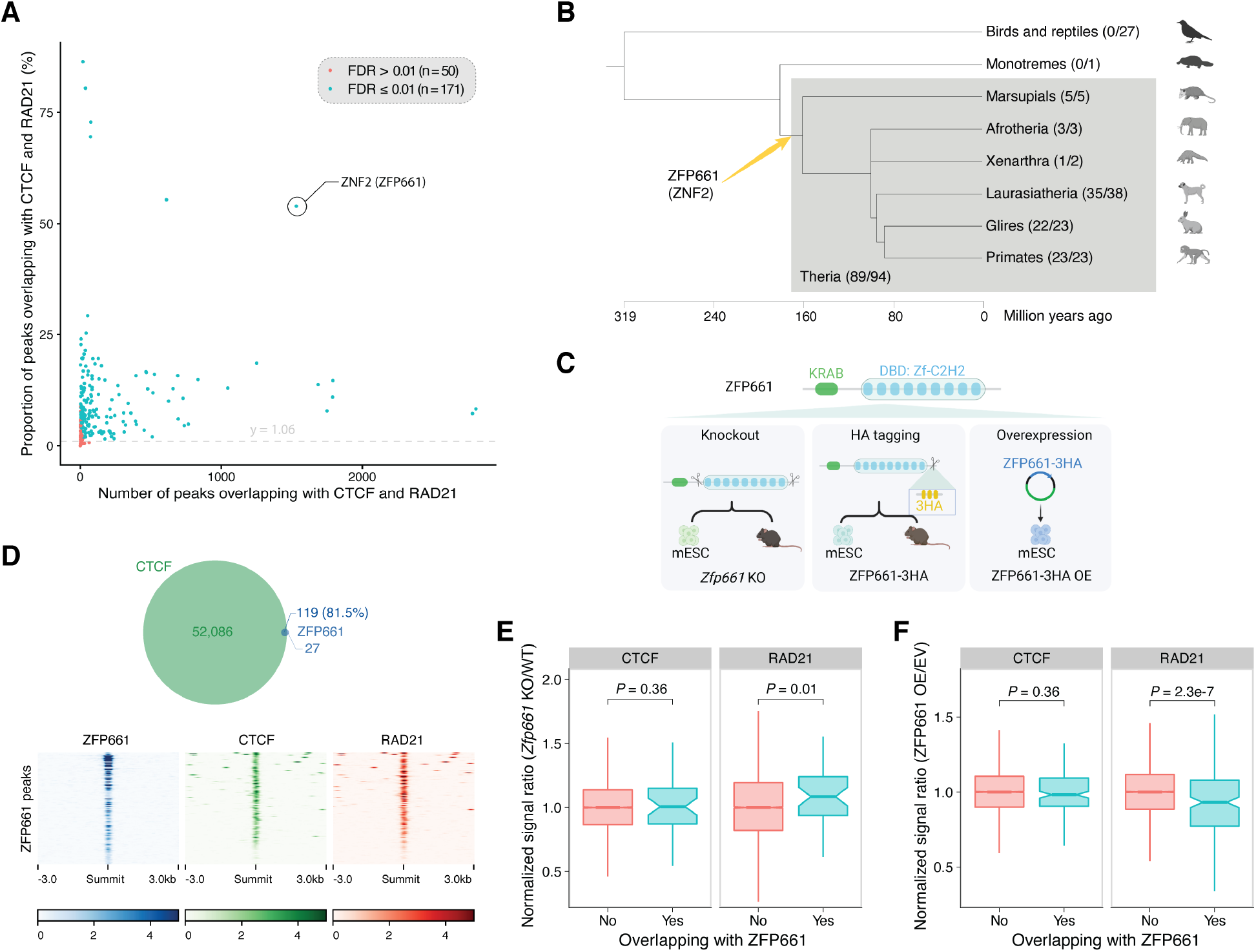
ZFP661, a conserved KRAB-ZFP in Theria, suppresses cohesin binding at CTCF sites without altering CTCF binding. (**A**) Overlap of binding peaks for 221 human KRAB-ZFPs with those of CTCF and RAD21 for screening purposes. The proportion of random peaks overlapping with CTCF and RAD21 was used as background (y=1.06), and single-tailed exact binomial tests were performed (threshold: FDR adjusted *P* ≤ 0.01). (**B**) Presence/absence of ZFP661 in different vertebrate clades. ZFP661orthologs are detected in most therian genomes (89/94) but not monotremes or birds/reptiles. (**C**) Cell and mouse models generated to explore ZFP661 function. (**D**) Overlap of ZFP661 binding peaks with CTCF and RAD21 peaks in ZFP661-3HA mESCs. (**E and F**) Influence of ZFP661 on CTCF and RAD21 binding in *Zfp661* knockout mESCs (E) and ZFP661-3HA OE mESCs (F). CTCF and RAD21 peaks that do not overlap with ZFP661 peaks were used as controls. ZFP661 binding significantly suppresses the binding of RAD21 but not CTCF in both *Zfp661* knockout mESCs (E) and ZFP661-3HA OE mESCs (F). EV represents empty vector control. Single-tailed Wilcoxon rank sum tests were performed.

In mice, *Zfp661* mRNA is broadly expressed at low levels with detectable expression in mouse embryonic stem cells (mESCs) and relatively higher expression in the developing brain compared to other organs (fig. S2). To explore the function of ZFP661 during embryonic development, we generated *Zfp661* knockout (KO) mESCs and mice, endogenously tagged ZFP661-3HA mESCs and transgenic mice using CRISPR-Cas9 technology, and ZFP661-3HA stably overexpressing (OE) mESCs using lentiviral vectors (Fig. 1C). We determined ZFP661 binding locations in the mouse genome using ChIP-seq, revealing that over 80% of ZFP661 binding peaks overlapped with those of CTCF and cohesin in both endogenously tagged ZFP661-3HA mESCs (Fig. 1D) and ZFP661-3HA OE mESCs (fig. S3), consistent with its ortholog ZNF2 in human (Fig. 1A). We further identified a putative ZFP661 binding motif that closely matched a computationally predicted ZFP661 binding motif, which was present in all 200 peak regions that were used to detect motifs (fig. S4).

Unlike many KRAB-ZFPs that strongly recruit the co-factor KAP1 and subsequently SETDB1 to establish H3K9me3 heterochromatin at genomic targets (*28, 31, 32*), ZFP661 target regions only presented weak KAP1 and H3K9me3 signals after overexpression of ZFP661 (fig. S5), indicating a potentially atypical functional mechanism. To determine whether ZFP661 might influence CTCF or cohesin binding, we performed ChIP-seq for CTCF and RAD21 in *Zfp661*^-/-^ mESCs and ZFP661-3HA OE mESCs relative to wild-type controls. Surprisingly, we found that ZFP661 decouples CTCF and cohesin binding; *Zfp661* loss-of-function increased RAD21 binding, whereas overexpression of ZFP661 suppressed RAD21 binding at CTCF sites co-occupied with ZFP661, without altering CTCF binding (Figs. 1E and 1F).

### ZFP661 binds exclusively inside CTCF loop anchors and allows cohesin to pass through CTCF barriers

To determine whether ZFP661 co-occupies targets simultaneously with CTCF, we performed ChIP-reChIP-seq assays with anti-HA (1^st^ ChIP) and anti-CTCF (2^nd^ ChIP) antibodies in ZFP661-3HA OE mESCs. These experiments confirmed that ZFP661 and CTCF are co-bound (Fig. 2A). Previous studies have demonstrated that chromatin loops occur primarily between two convergent CTCF binding sites (as shown in the upper portion of Fig. 2B) (*33, 34*) and that reversal of the orientation of CTCF binding sites prevents cohesin from being trapped at CTCF barriers (*16*). To gain insights into how ZFP661 might influence cohesin binding, we analyzed ZFP661 binding locations, indicated by ChIP summits, relative to CTCF barriers. We found that ZFP661 bound exclusively inside CTCF loop anchors, like cohesin (Fig. 2B). We confirmed this finding by analyzing the position of ZFP661 binding sites relative to CTCF binding sites, which were consistently located inside the loop anchors (fig. S6).

**Fig. 2.**
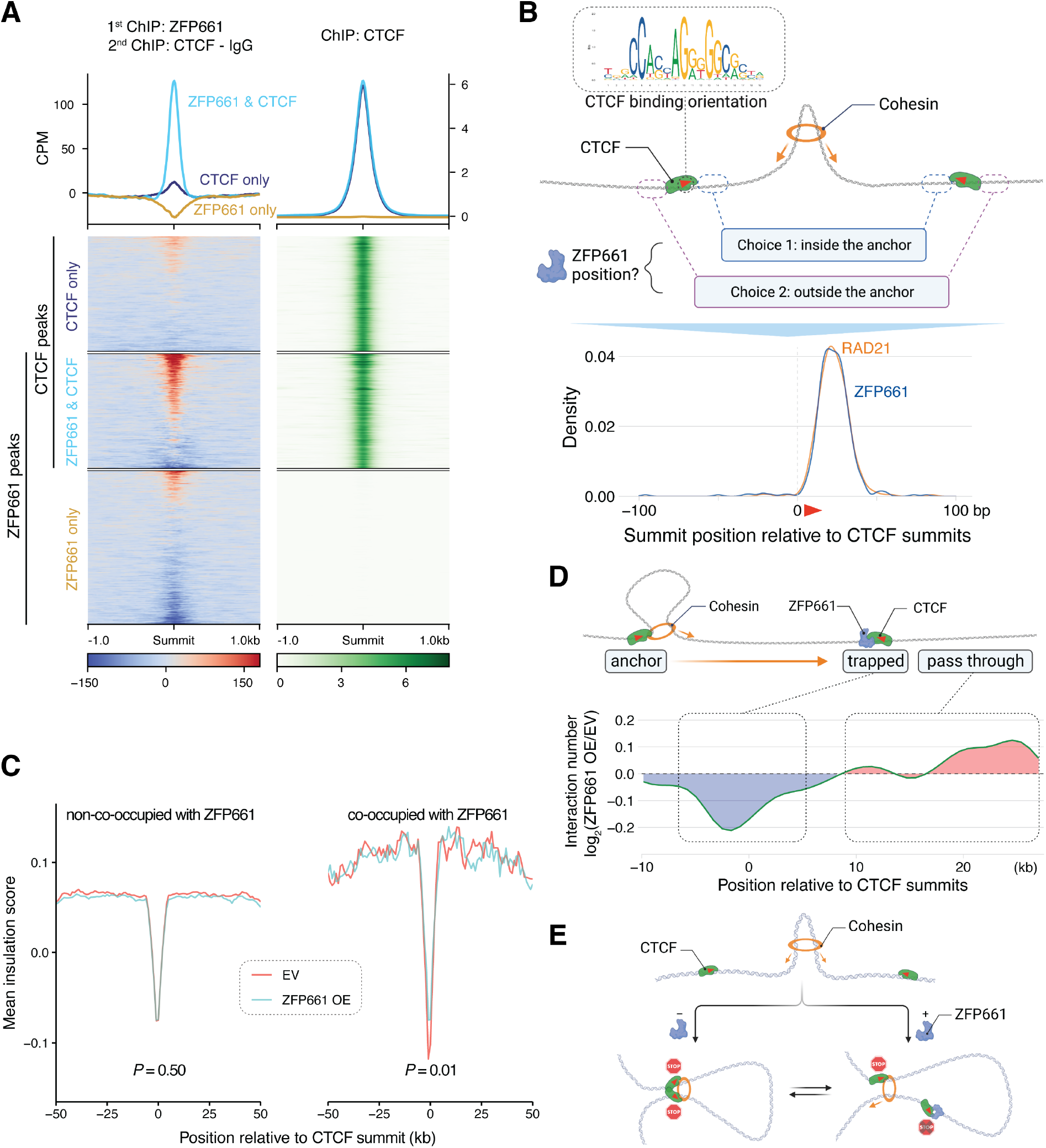
ZFP661 binds exclusively inside CTCF loop anchors and allows cohesin to pass through CTCF barriers. (**A**) ChIP-reChIP assay in ZFP661 OE mESCs shows that ZFP661 and CTCF simultaneously co-bind to targets. The 1^st^ ChIP was performed using anti-HA antibody, and the 2^nd^ ChIP was performed using anti-CTCF and anti-IgG antibodies, respectively. ChIP-reChIP signals in the heatmap represent anti-CTCF signals subtracted by anti-IgG signals. ZFP661-only and CTCF-only peaks, which are not co-occupied with each other, were used as controls. (**B**) Binding locations of ZFP661 and RAD21, indicated by ChIP summits, relative to those of CTCF. CTCF binding orientation is shown as the motif in the upper portion. (**C**) Influence of ZFP661 binding on the insulation score at CTCF barriers that are co-occupied with ZFP661. CTCF barriers that are not co-occupied with ZFP661 were used as controls. Single-tailed paired t-tests were performed. (**D**) Comparison of the distribution of chromatin loops trapped around or passing over CTCF barriers that are co-occupied with ZFP661 between EV control and ZFP661 OE. (**E**) Schematic diagram illustrates the working model of ZFP661. ZFP661 binding at CTCF barriers blocks cohesin trapping and allows cohesin to pass through the barriers.

We hypothesized two possible mechanisms that could explain ZFP661 decoupling of CTCF and cohesin signals: 1) ZFP661 could allow cohesin to pass through CTCF barriers or 2) ZFP661 could promote cohesin release from chromatin. To distinguish amongst these hypotheses, we tested the insulation scores at CTCF barriers and the proportion of chromatin loops that passed through CTCF barriers, reasoning that cohesin release would not decrease the barrier function of CTCF. We generated high-resolution Hi-C maps (1.18 billion valid interaction pairs in total) in ZFP661-3HA OE relative to empty vector (EV) controls. This allowed us to calculate both the insulation scores at CTCF barriers and the distribution of loop anchors at and over CTCF barriers. We found that overexpression of ZFP661 reduced insulation function of CTCF barriers (*i*.*e*., increased insulation scores) (Fig. 2C) and promoted loops to pass through CTCF barriers (rather than being trapped by CTCF) (Fig. 2D) at CTCF sites co-occupied with ZFP661.

Moreover, this effect was not seen at CTCF barriers that were not co-occupied with ZFP661 (Fig. 2C), demonstrating that ZFP661 binding suppresses the trapping of cohesin at CTCF barriers (Figs. 1E, 1F, and 2C). In sum, these results demonstrate that ZFP661 suppresses cohesin binding by allowing cohesin to pass through CTCF barriers. Considering that the interaction of the CTCF N-terminus (the portion facing cohesin extrusion and ZFP661 binding) with cohesin is critical for cohesin trapping (*35*), combined with our finding that ZFP661 binds at the same location where cohesin is typically trapped (Fig. 2B), we speculate that ZFP661 suppresses cohesin trapping by decreasing the probability of cohesin interaction with CTCF and/or competition with trapped cohesin (Fig. 2E).

### ZFP661 promotes the interaction of downstream enhancers with distal Pcdh promoters

To determine the potential role of ZFP661 during development by modulating the strength of CTCF barriers, we generated *Zfp661* null (*Zfp661*^-/-^) mice (Fig. 1C). *Zfp661*^-/-^ mice were viable and fertile, and displayed no overt phenotypes. Therefore, we suspected that ZFP661 might play a more subtle role in development. We performed Gene Ontology enrichment analysis of ZFP661 target genes (based on proximity to binding sites) and identified “homophilic cell adhesion via membrane adhesion molecules” as the most significantly enriched biological process (Fig. 3A). Interestingly, 21 of 25 target genes associated with this term belong to the protocadherin (Pcdh) gene clusters (the occupancy of ZFP661 and CTCF at the *Pcdh* locus is shown in Fig. 3B and fig. S7A). There are three Pcdh gene clusters (*Pcdhα/a, Pcdhβ/*b *and Pcdhγ/g*) in mammalian genomes, representing 14 *Pcdhα* and 21 *Pcdhγ* isoforms (excluding *Pcdhgb8*, which is annotated as a non-coding gene), each with alternative promoters, and 22 *Pcdhβ* genes in the mouse genome (fig. S7A). *Pcdhs* are activated by downstream enhancers dependent on the cooperation of CTCF and cohesin, where CTCF located at each promoter can stop cohesin extrusion to facilitate the contact of this promoter with downstream enhancers (*14-16*).

**Fig. 3.**
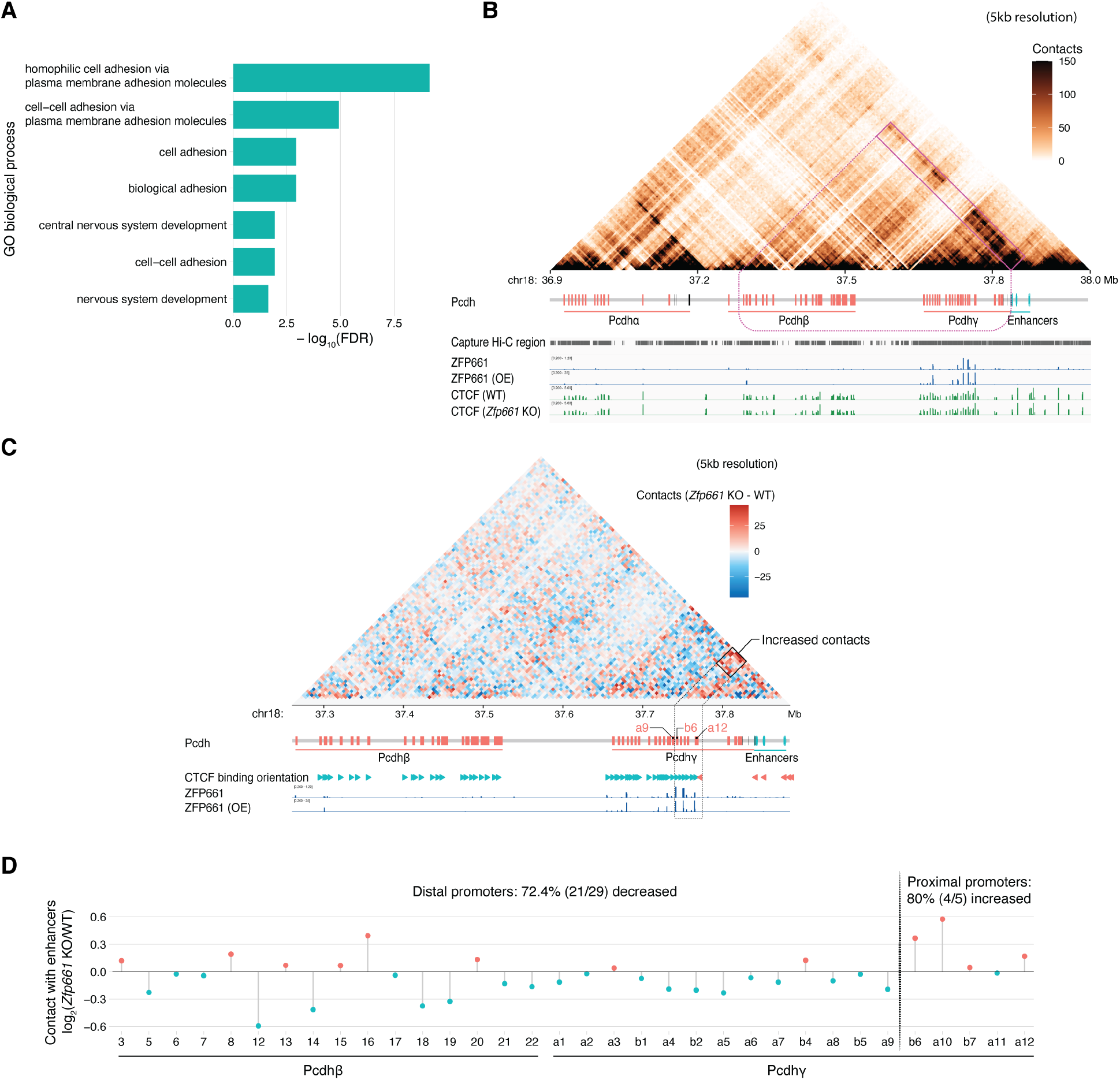
ZFP661 promotes the interaction of *Pcdhγ* downstream enhancers with distal protocadherin (Pcdh) promoters. (**A**) Gene Ontology (GO) enrichment analysis of ZFP661 target genes in “biological processes”. (**B**) Capture Hi-C map at the *Pcdh* locus in E16.5 mouse forebrains (upper) and ZFP661 and CTCF occupancy in mESCs (lower). The Capture Hi-C data demonstrate that the enhancer cluster downstream of *Pcdhγ* locus interacts with both *Pcdhβ* and *Pcdhγ* promoters. (**C and D**) Alterations in contacts at the *Pcdhβ* and *Pcdhγ* regions (C) and contacts between the enhancer cluster and individual promoters (TSS ± 2.5 kb) (D) after *Zfp661* knockout in the Capture Hi-C map of E16.5 forebrains. Loss of *Zfp661* leads to increased contacts of enhancers with proximal isoforms (C) and reduced contacts with distal promoters (D).

At the *Pcdh* locus, ZFP661 binding regions were co-bound by CTCF and were primarily located at the proximal promoters closest to the downstream enhancer cluster of *Pcdhγ* (Fig. 3B and fig. S7A). We generated capture Hi-C maps in mouse E16.5 forebrains (when neurons occupy the largest proportion by cell population in the developing mouse brain (*36*)) and demonstrated that the downstream enhancers of *Pcdhγ* interact with *Pcdhβ* and *Pcdhγ* clusters (Fig. 3B), indicating they control both *Pcdhβ* and *Pcdhγ* expression (*37*). We reasoned that ZFP661 binding at enhancer proximal promoters might promote cohesin to pass through CTCF barriers, allowing the enhancers to contact and activate more distal promoters of the *Pcdhβ* and *Pcdhγ* clusters. To test this hypothesis, we performed Capture Hi-C experiments of the *Pcdh* locus in E16.5 forebrains of *Zfp661*^+/+^ and *Zfp661*^-/-^ littermates. Importantly, we found that loss of *Zfp661* increased the contacts of enhancers with proximal promoter regions (4/5 increased) while decreasing contacts with more distal promoters (21/29 decreased; Figs. 3C and 3D). Consistent with this result, we found that ZFP661 overexpression (in ZFP661-3HA OE mESCs) decreased the contacts of these enhancers with proximal regions (fig. S7B) and promoted contacts with distal promoters (fig. S7C).

### ZFP661 balances the usage of Pcdh isoforms and increases Pcdh expression diversity

Previous studies have demonstrated that the interaction between downstream enhancers and *Pcdh* alternative promoters is the basis for activating *Pcdh* expression (*15, 16, 37*). To investigate how the alteration of enhancer-promoter interactions affects the *Pcdh* expression repertoire in individual neurons, we performed single-cell 5’ RNA-seq in mouse E16.5 forebrains to distinguish each Pcdh isoform. The number of unique molecular identifiers (UMIs) and genes detected per cell between *Zfp661*^+/+^ and *Zfp661*^-/-^ littermates were matched by adjusting for sequencing depth and by down-sampling UMIs to avoid possible biases in downstream analyses (figs. S8A and S8B). Cell types were assigned by projection onto a cell atlas of developing mouse brains at the corresponding ages (*36*)(fig. S8C and Fig. 4A). Compared with *Zfp661*^+/+^ brains, *Zfp661*^-/-^ brains had no obvious alterations in cell populations (Fig. 4A) or in the overall expression level of *Pcdhs* within cortical or hippocampal glutamatergic neurons (referred to as “glutamatergic neurons”) or forebrain GABAergic neurons (referred to as “GABAergic neurons”) (Fig. 4B). Compared with GABAergic neurons, glutamatergic neurons expressed higher levels of *Pcdhs* (Fig. 4B), perhaps due to their requirement for farther neurite projection (*38*). When comparing the usage of Pcdh isoforms, *Zfp661*^-/-^ brains had greater usage of enhancer proximal isoforms at the expense of distal isoforms in pooled neurons (Fig. 4C), glutamatergic neurons (fig. S9A) and GABAergic neurons (fig. S9B), consistent with the underlying chromatin-interaction results (Figs. 3C and 3D).

**Fig. 4.**
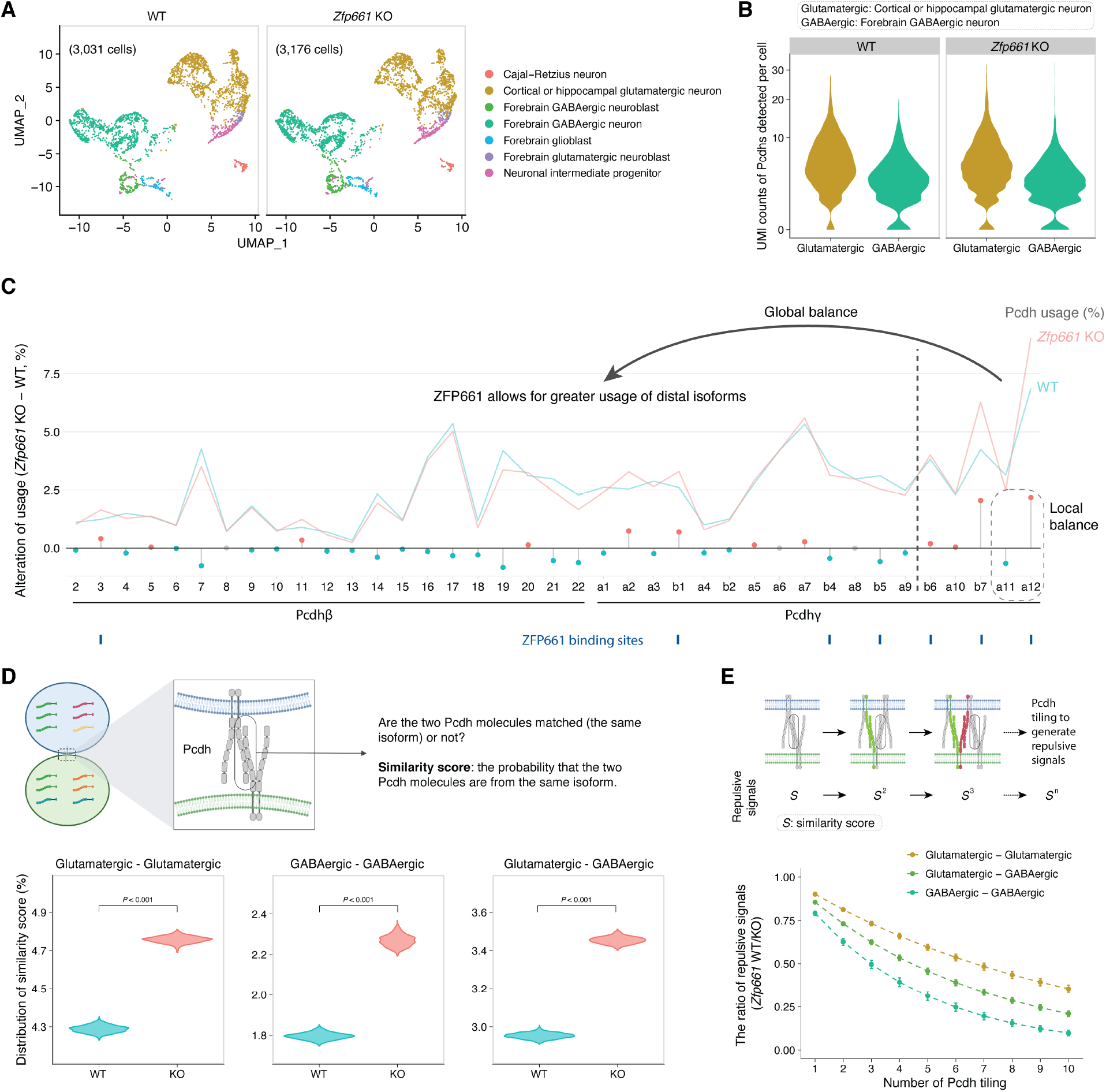
ZFP661 balances the usage of Pcdh isoforms and increases the diversity of Pcdh expression. (**A**) UMAP visualization of the cell atlas in *Zfp661*^+/+^ and *Zfp661*^-/-^ E16.5 mouse forebrains. (**B**) Unique molecular identifier (UMI) counts of Pcdhs detected in glutamatergic and GABAergic neurons of *Zfp661*^+/+^ and *Zfp661*^-/-^ brains. The y-axis was square-root transformed. (**C**) Usage alterations of Pcdhβ/γ isoforms in pooled glutamatergic and GABAergic neurons of *Zfp661*^+/+^ and *Zfp661*^-/-^ brains. The lines represent isoform usage, and lollipops represent the difference in isoform usage between *Zfp661*^+/+^ and *Zfp661*^-/-^ neurons. ZFP661 allows for greater usage of distal Pcdh isoforms. (**D**) The distribution of median similarity scores of the Pcdh repertoire between all pairs of single neurons within glutamatergic (left), GABAergic (center), or between glutamatergic and GABAergic neurons (right) in *Zfp661*^+/+^ and *Zfp661*^-/-^ brains. Similarity scores within each type were subsampled 50% of the datapoints for 1000 times, and *P* values were calculated based on the 1000 subsamples. (**E**) Ratio of repulsive signals between *Zfp661*^+/+^ and *Zfp661*^-/-^ neurons by simulating Pcdh chain formation. Repulsive signal represents the probability that a Pcdh isoform from one neuron matches the isoform from another neuron during continuous comparisons. ZFP661 decreases Pcdh-based repulsive signals amongst neurons (data are shown as mean ± SD).

Diversity in the Pcdh expression repertoire allows neurites to co-share local space and form more complex connections (*5*). To better quantify Pcdh diversity and explore the impact of Pcdh usage alteration in *Zfp661*^-/-^ neurons, we calculated a similarity score between all pairs of cells within the population, which represents the probability that a Pcdh isoform taken from one cell will match an isoform from another cell (as shown in the upper portion of Fig. 4D). By comparing median similarity scores of 1000 subsamples from all pairs of cells within each population, we determined that *Zfp661*^-/-^ neurons possessed significantly greater similarity (*i*.*e*., reduced diversity) than *Zfp661*^+/+^ neurons, both within glutamatergic and GABAergic neuron types and between the two types (Fig. 4D). To further investigate the impact of this reduced diversity on neuronal dendrite repulsion, we calculated repulsive signals, which represent the probability of sequential matching of Pcdh isoforms during continuous comparison by stimulating the chain formation of Pcdh molecules (*9*) (as shown in the upper portion of Fig. 4E). The results demonstrate that ZFP661 dramatically decreases repulsive signals from Pcdh chain formation, both within glutamatergic and GABAergic neuron types and between types (Fig. 4E). In sum, these data reveal that the loss of *Zfp661* leads to reduced complexity in Pcdh expression, which increases the likelihood that a cell will encounter another cell with the same or highly similar Pcdh expression repertoire. This subsequently results in increased repulsive signals during dendritic projection, potentially reducing the number of dendrites that can fit in a local space.

### Loss of *Zfp661* causes deficits in dendritic arborization and social interaction

To determine whether the reduced Pcdh expression diversity in *Zfp661*^-/-^ mice might cause dendritic projection defects, we performed Golgi-Cox staining on early adult mouse brains (P60-66) and imaged pyramidal neurons at layer II/III of somatomotor areas (Bregma: 1.70 to 1.18 mm), where we could easily ensure consistency in neuron type and location during sampling. Sholl analysis, which measures the complexity of neuronal dendrites, confirmed that *Zfp661*^-/-^ neurons have deficits in arborization and the distribution of neuronal dendrites (fig. S10 and Fig. 5A). Compared with *Zfp661*^+/+^ neurons, *Zfp661*^-/-^ neurons displayed a significant decrease in both the branch number (Fig. 5B) and total branch length of neuronal dendrites (Fig. 5C). These data are consistent with the hypothesis that the loss of *Zfp661*, which causes greater repulsive signals during Pcdh tiling (Fig. 4E), might inhibit the elongation and arborization of neuronal dendrites. This is also consistent with previous studies that demonstrated that direct decreases in Pcdh diversity (via genetic ablation) also cause deficits in dendritic arborization (*39-41*). Finally, we explored how *Zfp661* loss-of-function might impact behavior. Sociability tests revealed that *Zfp661*^-/-^ mice displayed deficits in social interactions (Fig. 5D), similar to mice that have reduced Pcdh diversity (*42, 43*).

**Fig. 5.**
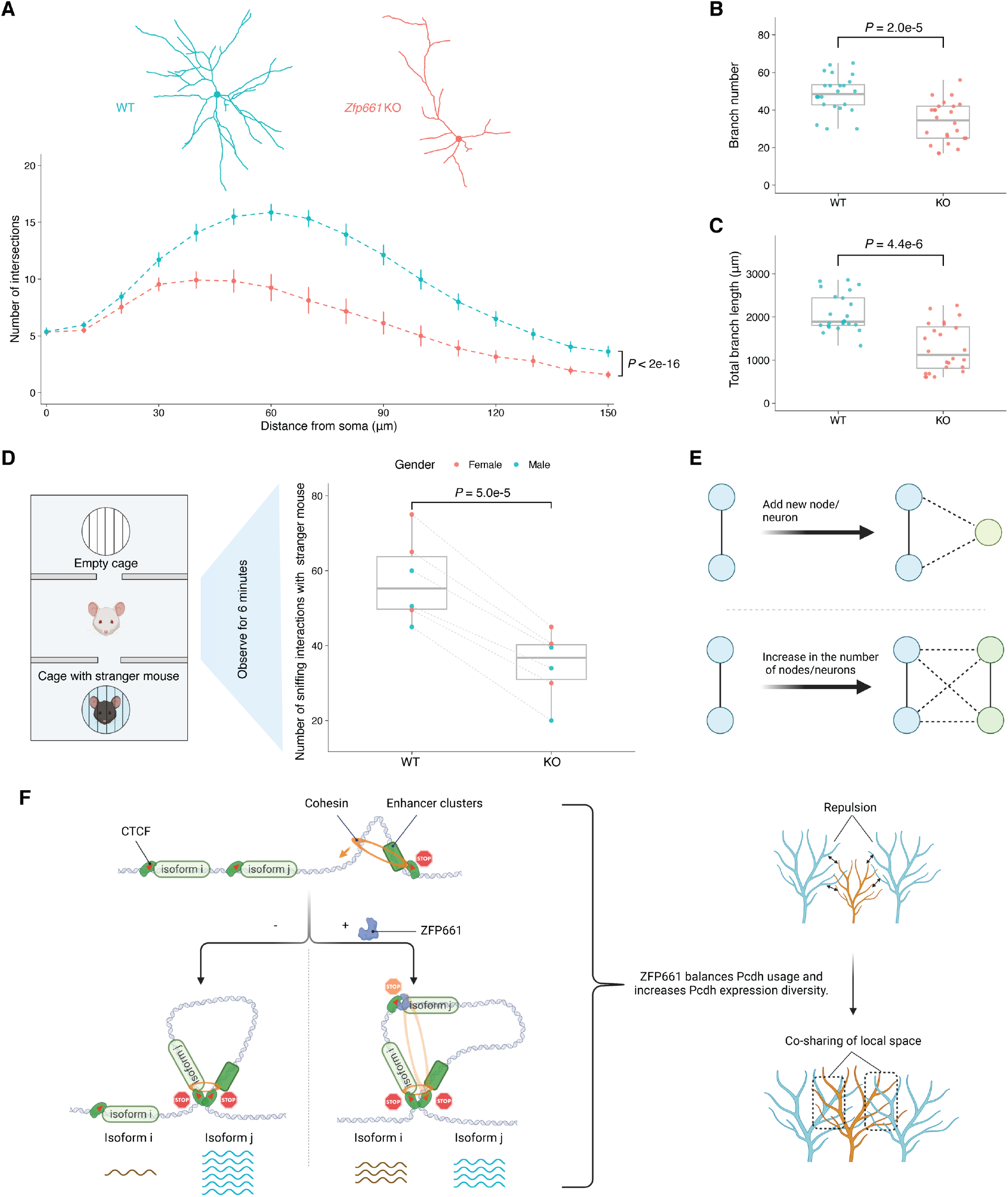
Loss of ZFP661 causes deficits in neuronal dendritic arborization and social interaction. (**A**) Sholl analysis of dendrites of pyramidal neurons in layer II/III of the cerebral cortex in *Zfp661*^+/+^ and *Zfp661*^-/-^ mouse brains (P60-66). Representative examples of 3D-reconstructions of *Zfp661*^+/+^ and *Zfp661*^-/-^ neurons are shown above (data are shown as mean ± SEM; n=24, 8 neurons per mouse and 3 mice per genotype, one-way repeated measures ANOVA).(**B and C**) Comparison of the branch number (B) and total branch length of neuronal dendrites (C) between *Zfp661*^+/+^ and *Zfp661*^-/-^ neurons (n=24, single-tailed Wilcoxon rank sum test). (**D**) *Zfp661*^-/-^ mice show social interaction deficits in sociability tests (n=6, 3 male and 3 female mice per genotype, single-tailed paired Wilcoxon rank sum test). Littermate pairs between *Zfp661*^+/+^ and *Zfp661*^-/-^ mice are connected using dashed lines. (**E**) Two simplified diagrams demonstrating the necessity of additional connections (dashed edges) for new neurons (green nodes) to integrate well into original brain networks to avoid fragmentation. (**F**) Schematic diagram illustrates that ZFP661 increases Pcdh expression diversity and promotes intermingling and co-sharing of local space amongst neuronal dendrites by balancing the expression of Pcdh isoforms. This may have helped to accommodate the larger mammalian brain.

## Discussion

CTCF binding is well known to act as a barrier for cohesin extrusion, but how CTCF stops the extrusion of cohesin is still unclear. Genetic modifications, including reversal of the orientation of CTCF binding sites (*16*) and destruction of CTCF-cohesin interaction (*35*) can suppress cohesin trapping without altering CTCF binding. These highlight the roles of CTCF binding orientation and CTCF-cohesin interactions in cohesin trapping. Here, we uncovered a novel mechanism that decouples CTCF and cohesin binding; ZFP661 exclusively binds inside CTCF loop anchors (Fig. 2B) and allows cohesin to pass through CTCF barriers (Figs. 2D and 3C and fig. S7B), indicating that the accessibility of CTCF-cohesin interface is crucial for cohesin trapping. The chromatin loops established by CTCF and cohesin confine over 90% of enhancer-promoter interactions that occur within CTCF boundaries (*21*). Consequently, the need for enhancer-promoter interactions to occur across specific CTCF barriers may require the breaking of CTCF boundaries. Our findings demonstrate that ZFP661 directly regulates CTCF-mediated cohesin trapping at CTCF barriers, providing a potentially rapid and reversible way to open/close CTCF boundaries.

During neural development, 3D genome structures are dramatically changed to accompany altered gene expression patterns (*44*). Numerous KRAB-ZFPs that originated during mammal evolution (*23, 25*) are preferentially expressed in developing brains (*29*) and bind at CTCF barriers (Fig. 1A), similar to ZFP661. Our findings suggest that KRAB-ZFP-dependent CTCF/cohesin modulation may have played important roles in driving the evolution of 3D genome structures in mammalian brain development.

Nodes and edges (*i*.*e*., connections amongst nodes) are two key components of a network. Increased connections per node are required after node expansion to maintain proper network architecture and avoid network fragmentation (as shown in the simplified cases in Fig. 5E), which also holds true in neuronal networks. During mammalian evolution, ZFP661 emerged as a factor that increases the diversity of Pcdh expression (Fig. 4D), which decreases the repulsive signals amongst neurons during dendritic projection (Fig. 4E). We hypothesize that this allows more neuronal dendrites to fit in a tighter space and promotes the arborization and elongation of neuronal dendrites during development (Fig. 5F), providing a possible solution that ensures proper neuronal connections and network preservation in the expanded mammalian brain.

Multiple findings indicate this function is critical throughout Theria: 1) ZFP661 is retained in most therian genomes (Fig. 1B); 2) ZFP661 has a highly conserved DNA binding domain with 97.2% (35/36) of fingerprint amino acids that determine DNA bases to contact being conserved (fig. S1); 3) the ZFP661/ZNF2 binding motif (fig. S11A), its association with CTCF binding (Figs. 1A and 1D), and its binding at the *Pcdh* locus (fig. S11B) are conserved between human and mouse.

Both the loss of *Zfp661* (Figs. 5A to 5D) and reduced Pcdh diversity from direct genetic ablation (*39-43*) can induce cortical dendritic arborization defects and social deficits. Autism Spectrum Disorder (ASD) is one of most common brain developmental disorders in humans associated with social deficits, which impacts about 2.3% children in the United States (*45*). The causes of ASD are largely unclear, but growing evidence suggests that defects in neuronal connections are a major underlying cause (*46*). Remarkably, ZNF2 binding at the *Pcdh* locus is conserved in humans (fig. S11B), and structural variations at *ZNF2* (fig. S12) and *Pcdh* loci (*47*) are associated with ASD in humans. These results highlight an important link between the regulation of Pcdh diversity by ZFP661(ZNF2), neuronal connections, and ASD.

## Acknowledgments

We thank A. Grinberg, J. Yimdjo and V. Biggs for generating and maintaining transgenic mice; T. Li, J. Iben, R. Lazris, F. Faucz and S. Coon for next-generation sequencing (NGS) supports; H. Potts and M. Peiravi for mouse behavioral tests; members of Macfarlan lab, K. Pfeifer, J. Kassis and P. Rocha for helpful discussion; J. Kassis, B. Shen, M. Bruno, M. Custance and G. Tricola for critically reading the manuscript and helpful suggestions. Most of the computational analyses were performed on the NIH Biowulf high-performance computing platform.

## Author contributions

Conceptualization: JJ, TSM

Methodology: JJ, TSM

Investigation: JJ, SR, EW, GW, DAS

Software: JJ

Formal analysis: JJ, MAS

Visualization: JJ

Funding acquisition: TSM

Project administration: TSM

Supervision: TSM

Writing – original draft: JJ

Writing – review & editing: JJ, TSM, RLC, ADS

## Data and materials availability

NGS data are available at Gene Expression Omnibus (GSE231869). Raw imaging data are available upon request, and preprocessed imaging data and neuronal traces are available at Mendeley (*48*). Program codes for calculating similarity scores for Pcdhs amongst neurons are available at Zenodo (*49*). Cell and mouse models generated in this work are available under material transfer agreements with National Institutes of Health.

## Supplementary Materials

Materials and Methods

Figs. S1 to S12

Tables S1 to S3

