## Supplementary Materials for "CTCF barrier breaking by ZFP661 promotes protocadherin diversity in mammalian brains"

### Materials and Methods

#### Cell culture

The following cell lines were used in this study: R1 mouse embryonic stem cells (mESCs) (129S1/SvImJ x 129X1/SvJ background), JM8.F6 mESCs (C57BL/6N background), and Lenti-X 293T cells (Takara, 632180). The mESCs were cultured in ES medium, which consisted of DMEM (Gibco, 11995-065), 15% FBS (EmbryoMax, ES-009-B), 10mM HEPES (Gibco, 15630-080), 1X MEM Non-Essential Amino Acids Solution (Gibco, 11140-050), 1X GlutaMax Supplement (Gibco, 35050-061), 1X 2-Mercaptoethanol (Gibco, 21985-023), 1000 units/ml Recombinant mouse LIF (ESGRO, ESG1107), and 1X Antibiotic-Antimycotic (Gibco, 15240-062). mESCs were thawed onto mitomycin C-treated MEF feeder cells (EmbryoMax, PMEF-N) and subsequently passaged onto dishes coated with 0.1% Gelatin Solution (EmbryoMax, ES-006-B). Lenti-X 293T cells were cultured in DMEM medium supplemented with 10% FBS. All cells were cultured in a 37 °C humidified incubator with 5% $CO_2$. Mycoplasma tests were routinely performed using the Mycoplasma PCR Detection Kit (abm, G238), and the cell lines used were negative for Mycoplasma contamination.

#### Analysis of public ChIP-seq data for human KRAB-ZFPs

The raw ChIP-seq data for 221 human KRAB-ZFPs (tagged with HA and overexpressed in HEK293T cells) and ChIP-seq input were downloaded from GEO (GSE78099) (*1*). Raw fastq reads were cleaned using Trimmomatic (v0.38) (*2*) to remove sequencing adaptors and low-quality bases from both ends with the parameters "LEADING:3 TRAILING:3 SLIDINGWINDOW:4:15 MINLEN:25". The resulting clean reads were mapped onto the human genome (hg19) using BWA (*3*) with default parameters, and only uniquely mapped reads (with tag X0:i:1) were retained for downstream analyses. MACS2 (v2.2.7.1) (*4*) was used to call peaks with ChIP input as a control, one duplicated tag kept, the fragment length estimated, and the threshold $q \leq 0.01$. Peaks that overlapped (overlap represents any bases of a peak sharing with other peaks/regions) with the regions on ENCODE blacklist (*5*) were filtered out. The annotation of CTCF and RAD21 binding peaks was obtained from ENCODE (v2, cell type: H1-hESC) (*6*). The proportion of random peaks (with the same average length as KRAB-ZFPs peaks) overlapping with CTCF and RAD21 co-occupied peaks was calculated (1.06%) and used as the background. Single-tailed exact binomial tests were performed to determine whether each KRAB-ZFP presented more peaks that overlapped with CTCF and RAD21 co-occupied peaks than the random background, and the resulting *P* values were adjusted using FDR.

#### ZFP661/ZNF2 conservation analyses

The presence/absence of ZFP661/ZNF2 in vertebrate genomes was determined based on the "gene gain/loss tree" in Ensembl (v107) (*7*), and divergent times amongst different clades were obtained from TimeTree (v5) (*8*). The multiple sequence alignment of ZNF2/ZFP661 was performed using CLUSTAL Omega (v1.2.4) (*9*), and annotations of KRAB domain and C2H2 zinc-finger arrays were obtained from UniProt (*10*). The fingerprint amino acids were labelled based on the relative positions within zinc-finger arrays according to the reference (*11*).

#### Generation of transgenic cell lines and mouse models using CRISPR-Cas9 technology

Endogenously tagged ZFP661-3HA mESCs and *Zfp661* knockout (KO) mESCs were generated using CRISPR-Cas9 technology. To generate endogenously tagged ZFP661-3HA mESCs, JM8.F6 mESCs (C57BL/6N background) were used. One guide RNA (gRNA) (the guide sequence was shown in table S1) was designed to target close to the Zfp661 stop codon, and single-stranded oligodeoxynucleotides (ssODNs) (the sequence was shown in table S2) were used as a repair template to insert a 3xHA tag at the C-terminus of ZFP661. To generate *Zfp661* KO mESCs, R1 ESCs (129S1/SvImJ x 129X1/SvJ background) were used. Two gRNAs (table S1) were designed to target both sides of the ZFP661 DNA-binding domain and delete the DNA-binding domain. Briefly, guide sequences (table S1) were cloned into pX330-U6-Chimeric_BB-CBh-hSpCas9 plasmid (Addgene, #42230) at the BbsI cutting site. 10 µg of pX330 plasmid with guide sequences (for *Zfp661* KO, 5 µg of each plasmid) and 1 µg of pPGKpuro plasmid (Addgene, #11349) were co-transfected into $2x10^6$ mESCs using the Nucleofector-II (program A-023). The cells were then plated onto a 10cm dish with mitomycin C-treated puromycin-resistant MEF feeder cells and replaced with fresh ES medium containing 1 µg/ml puromycin (Gibco, A11139-03) 24 hours post-nucleofection. Single colonies were picked about one-week post-nucleofection and genotyped using PCR with the primers listed in table S3. The designated positive colonies were further confirmed using Sanger sequencing. Two biological replicates for each condition (*Zfp661*$^{+/+}$ and *Zfp661*$^{-/-}$) were obtained from two different single colonies picked, respectively.

To generate endogenously tagged ZFP661-3HA transgenic mice, mESCs from one heterozygous cell line were injected into blastocysts from B6 albino mice (Jax, 000058) following the standard protocol. The resulting transgenic carrier was further crossed with C57BL/6J mice (Jax, 000664). The background of maintained ZFP661-3HA transgenic mice is a mixture of C57BL/6 and B6 albino. The *Zfp661* KO mice were generated using zygote microinjection. Two synthetic sgRNAs (Synthego) with the same guide sequences used in generating *Zfp661* KO mESCs (table S1) and SpCas9-2NLS nuclease (Synthego) were assembled into RNP complexes at a ratio of 1:1 and injected into zygotes obtained from B6D2F1/J mice (Jax, 100006) following the Synthego mouse embryo microinjection protocol. The resulting transgenic carrier was further crossed with C57BL/6J mice. The background of *Zfp661* KO mice maintained is a mixture of C57BL/6J and B6D2F1/J.

Mice were housed with a light cycle of 14-hour on/10-hour off, temperature maintained between 20 to 22°C, humidity kept at 40-55%, and with no more than 5 mice per cage. All mouse procedures had been reviewed and approved by the NICHD ACUC at the National Institutes of Health (ASP#: 18-026). Mice were euthanized using $CO_2$ following the approved procedure. Only *Zfp661*$^{+/+}$ and *Zfp661*$^{-/-}$ littermate pairs were used in this study.

Generation of ZFP661-3HA overexpressing mESCs using lentiviral vectors

The DNA sequence of ZFP661-3HA was synthesized with codon usage optimization and cloned into the pLV-EF1a-IRES-Blast plasmid (Addgene, #85133) between the BamHI and EcorI cutting sites, resulting in the ZFP661-85133 plasmid. To package the lentivirus, 4 µg of ZFP661-85133 plasmid (or pLV-EF1a-IRES-Blast plasmid for empty vector (EV) control), 4 µg of pMDLg/pRRE plasmid (Addgene, #12251), 2 µg of pRSV-Rev plasmid (Addgene, #12253) and 2 µg of pMD2.G plasmid (Addgene, #12259) were co-transfected into Lenti-X 293T cells cultured in a 10cm dish. The medium was replaced with fresh medium 24 hours post-transfection, and the lentivirus-containing supernatant was harvested 48 hours post-transfection. Debris was removed by centrifugation at 1000 g for 10 min.

To transduce R1 ESCs, 50,000 cells were seeded onto 6-well plates with ES media containing 8 μg/ml polybrene (Millipore, TR-1003-G) and 100 ul of lentivirus-containing supernatant (lenti-EV or lenti-ZFP661). Selection was performed 48 hours post-transduction by adding fresh ES medium containing 6 μg/ml Blasticidin S HCl (Gibco, A11139-03), which was continued for 10 days to remove all negative cells. Two biological replicates for each condition (EV control and ZFP661-3HA OE) were obtained from two independent transductions, respectively.

ChIP-seq in mESCs and data processing

Chromatin immunoprecipitation (ChIP) assays were performed according to the ENCODE protocol (*6*) with modifications. 20M cells/ChIP were used for ZFP661-3HA (anti-HA: Abcam, ab9110), CTCF (anti-CTCF: Millipore, 07-729), RAD21 (Anti-RAD21: Abcam, ab992) and KAP1 (anti-KAP1: Abcam, ab22553), and 10M cells/ChIP were used for H3K9me3 (anti-H3K9me3: Abcam, ab8898). 5 μg of antibodies were used for each ChIP. Briefly, 1) Cells were crosslinked with 1% formaldehyde (Thermo Scientific, 28908) for 10 min and quenched with 0.4M glycine for 5 min; 2) Crosslinked cells were lysed using 2.5 ml cell lysis buffer (5 mM PIPES pH 8.0, 85 mM KCl, 0.5% IGEPAL CA-630) with 1X fresh protease inhibitor cocktail (PIC) (Roche, 11836145001), and nuclei were collected and resuspended in 1 ml RIPA buffer (50 mM Tris-HCl pH 8.0, 150 mM NaCl, 2 mM EDTA pH 8.0, 1% IGEPAL CA-630, 0.5% Sodium Deoxycholate, 0.1% SDS) with 1X fresh PIC; 3) The resulting chromatin were sonicated to a fragment size of 200 to 500 bp using a Bioruptor Plus; 4) Dynabeads Protein A/G were blocked using 0.5% BSA/PBS blocking buffer (0.9 ml, 5 min x 3 times at 4°C). Chromatin fragments were precleared using 30 μl of Dynabeads Protein A/G for ~2 hours at 4°C, and 5 μg of antibodies were coupled to 30 μl of Dynabeads Protein A (Invitrogen, 10001D) for antibodies from rabbit or Dynabeads Protein G (Invitrogen, 10003D) for antibodies from mouse for 5-6 hours at 4°C and then washed using 0.5% BSA/PBS blocking buffer (0.9 ml, 5 min x 3 times at 4°C); 5) Beads with coupled antibodies were added to precleared chromatin fragments for immunoprecipitation (IP) overnight at 4°C; 6) Beads were washed using low salt wash buffer (0.1% SDS, 1% Triton X-100, 2 mM EDTA, 20 mM Tris-HCl pH 8.0, 150 mM NaCl; 5 min at 4°C), high salt wash buffer (0.1% SDS, 1% Triton X-100, 2 mM EDTA pH 8.0, 20 mM Tris-HCl pH 8.0, 500 mM NaCl; 5 min x 2 times at 4°C), LiCl wash buffer (250 mM LiCl, 1% IGEPAL CA-630, 1% Sodium Deoxycholate, 1 mM EDTA, 10 mM Tris-HCl pH 8.0; 5 min x 2 times at 4°C) and TE buffer (10 mM Tris-HCl pH 8.0, 1 mM EDTA; 5 min x 2 times at 4°C); 7) After washing, beads were resuspended in 200 μl of freshly made IP elution buffer (10 mM Tris-HCl pH8.0, 0.3 M NaCl, 5 mM EDTA pH 8.0, 0.5% SDS), added 2 μl of RNase A (10 mg/ml) and then incubated overnight at 65°C to reverse crosslinking. Then 3μl of Proteinase K (20 mg/ml) was added and the mixture was further incubated for 2 hours at 55°C. ChIP DNA was further purified using Genomic DNA Clean & Concentrator-10 (Zymo, D4011), and 10 μl (or up to 50 ng) of ChIP DNA was used as input to construct ChIP library using ThruPLEX DNA-Seq Kit (Takara, R400676). The ChIP inputs from the same biological replicate were evenly sampled and pooled together to construct the corresponding ChIP input library. Size selection (250bp to 600bp) was performed on the resulting libraries using HighPrep beads (MagBio, AC-60050). Libraries for *Zfp661* KO mESCs and endogenously tagged ZFP661-3HA mESCs were sequenced using HiSeq 2500 (single-end 50 bp), and libraries for ZFP661-3HA OE mESCs were sequenced using NovaSeq 6000 (paired-end 2x50 bp).

Using the same setting described in the "Analysis of public ChIP-seq data for human KRAB-ZFPs" section, Raw fastq reads were cleaned using Trimmomatic, and were mapped onto the mouse genome (mm10) using BWA. Only uniquely mapped reads were retained for downstream analyses. ChIP binding peaks were detected using MACS2 (with the fragment length 200 bp for single-end datasets) and were further filtered using the ENCODE blacklist. During peak calling, ChIP inputs were used as controls for all factors except for ZFP661. For ZFP661 in ZFP661-3HA OE mESCs, ChIP input and the corresponding ChIP in EV control mESCs using anti-HA antibodies (referred to as HA-ChIP) were used as controls to detect peaks (referred to as input-peaks and HA-peaks), respectively. Only HA-peaks that overlapped with input-peaks were retained as reliable peaks for downstream analyses. For ZFP661 in endogenous tagged ZFP661-3HA mESCs, HA-ChIP in un-tagged mESCs were used as ChIP input to detect peaks. For H3K9me3, the peak regions from MACS2 output were used directly. For other factors, the 50 bp flanking regions of ChIP summits (*i.e.*, summits ± 50 bp) were used as refined peak regions for downstream analyses.

To generate high-quality and consistent ChIP peak lists, peaks from two biological replicates in the same condition (*i.e.*, *Zfp661*$^{+/+}$ or *Zfp661*$^{-/-}$, EV control or ZFP661-3HA OE) were combined by retaining peaks in replicate 1 that overlapped with peaks in replicate 2. As ZFP661 expressed at low level in mESCs, the following steps were employed to generate a high-quality peak list for ZFP661 in ZFP661-3HA mESC: ZFP661 peaks in ZFP661-3HA mESCs were detected using a relatively loose threshold ($q \leq 0.1$), and only peaks that overlapped with consistent ZFP661 peaks in ZFP661-3HA OE mESCs were retained. To generate consistent ChIP signal (*i.e.*, bigwig) files, deepTools (v3.5.1) (*12*) was used with the parameters "--normalizeUsing CPM --binSize 10 –extendReads" (CPM represents counts per million) and with chrM and regions on ENCODE blacklist excluded. The ChIP signals were normalized by subtracting the ChIP input in each biological replicate (for ZFP661, subtracted by HA-ChIP), and then the two biological replicates were averaged to generate a consistent bigwig file for each factor.

Normalization of peak intensities for cross-condition comparison

During exploring the influence of ZFP661 binding on CTCF and RAD21 binding, CTCF and RAD21 peak intensities were normalized as follows: 1) the intensities of peaks (indicated by the 5$^{th}$ column in MACS2 peak/summit outputs) from two biological replicates were averaged; and 2) a scale number was calculated to make the medians of peak intensities equal for peaks that did not overlap with ZFP661 peaks in the control and treatment, and peak intensities were then normalized between *Zfp661*$^{+/+}$ and *Zfp661*$^{-/-}$, and between EV control and ZFP661-3HA OE by multiplying them with the corresponding scale numbers. The ratio of normalized intensities for CTCF and RAD21 peaks between *Zfp661*$^{-/-}$ and *Zfp661*$^{+/+}$, and between ZFP661-3HA OE and EV control were calculated, respectively. To avoid the effects of large noise when the intensities of CTCF and RAD21 peaks were low and insensitive to alterations when they were too high, only the interquartile (*i.e.,* middle 50%) peaks (ranked by the peak intensities) were used in the calculation of the signal ratio. Single-tailed Wilcoxon rank-sum tests were performed to determine whether CTCF and RAD21 bindings were suppressed by ZFP661 binding, respectively.

In this study, peaks non-overlapping with ZFP661 refers to peaks that did not overlap with any of the ZFP661 peaks in any replicate, using either HA-ChIP or ChIP input as a control. Without a specific statement, only the top 500 ZFP661 peaks (ranked by peak intensities) in ZFP661-3HA OE mESCs (covered 89.7% ZFP661 binding peaks detected in endogenous tagged ZFP661-3HA mESCs; fig. S3A) were used in this study.

Detection of ZFP661 binding motif

Using the top 200 ZFP661 peaks that did not overlap with CTCF peaks in ZFP661-3HA OE mESCs as input, MEME-ChIP (*13*) was used to identify ZFP661 binding motifs within the 50 bp flanking regions of the ChIP summits. In addition, a ZFP661 binding motif was also predicted only using its protein sequence with the "Expanded linear SVM" model selected and all nine zinc-fingers included (at website http://zf.princeton.edu/) (*14*). To compare whether the motifs derived from ChIP-seq data was similar to the *de novo* predicted motif, TomTom (*15*) was used with default parameters, and a gap was manually added to the motif derived from ChIP-seq data for a better alignment.

ChIP-reChIP-seq and data analyses

ChIP-reChIP was performed on two biological replicates of ZFP661-3HA OE mESCs. For each replicate, 40M cells were used and divided into two ChIP assays for the 1st ChIP. In the 1st ChIP, 5 µg of anti-HA antibodies per ChIP were used by following the ChIP protocol described in the "ChIP-seq in mESCs and data processing" section until finishing the step 6. Then, the beads of two ChIP assays from the same biological replicate were pooled and resuspended in 50 µl of re-ChIP elution buffer (10 mM Tris-HCl pH 8.0, 1 mM EDTA, 2% SDS, and 15 mM DTT) with 1X fresh PIC, and incubated 30 min at 37°C to elute chromatin and denature antibodies. The beads were then removed, and the chromatin supernatant was diluted to normal IP conditions by adding 950 µl of re-ChIP dilution buffer (0.01% SDS, 1.1% Triton X-100, 1.2 mM EDTA, 16.7 mM Tris–HCl pH 8.0, and 167 mM NaCl) with 1X fresh PIC. To remove any remnant anti-HA antibodies, chromatin was further precleared with 25 µl of Dynabeads Protein A by incubation for 2 hours, while 2.5 µg of anti-CTCF and anti-IgG antibodies (Invitrogen, 02-6102) were coupled onto 15 µl of Dynabeads Protein A, respectively. Then, chromatin from each biological replicate was divided into two parts (one for anti-CTCF and the other one for anti-IgG), and ChIP procedure from steps 5 to 7 were performed for the 2nd ChIP. ChIP DNA was purified, and library construction and size selection were performed as described in the "ChIP-seq in mESCs and data processing" section. The resulting libraries were sequenced using NovaSeq 6000 (paired-end 2 x 50 bp).

The raw fastq reads from the 2nd ChIP (anti-CTCF and anti-IgG in two biological replicates) were cleaned and mapped onto the mouse genome (mm10) using the same settings as described in the "Analysis of public ChIP-seq data for human KRAB-ZFPs" section. Only uniquely mapped reads were retained for downstream analyses. The top 1000 CTCF-only peaks (CTCF peaks that did not overlap with ZFP661 peaks), the top 1000 ZFP661&CTCF peaks (ZFP661 peaks that were co-occupied with CTCF peaks), and all ZFP661-only peaks (ZFP661 peaks that did not overlap with CTCF peaks, n = 1337) were used for following analysis. Uniquely mapped reads in the 1kb flanking regions of the ChIP summits of these peaks were extracted, and duplicated reads were removed using Samtools (*16*). The resulting reads from the two biological replicates were pooled together. DeepTools was used to generate the "CTCF - IgG" bigwig file (anti-CTCF signals subtracted by anti-IgG signals) with parameters "-normalizeUsing CPM --binSize 10 –extendReads" and excluding chrM and the regions on ENCODE blacklist. If

ZFP661 does not co-bind targets with CTCF simultaneously, the profile of "CTCF – IgG" signals on ZFP661&CTCF peaks would be similar to the overlay of the two profiles from CTCF-only peaks and ZFP661-only peaks.

Location of ZFP661 binding relative to CTCF binding

The CTCF binding motif (MA0139.1) was obtained from JASPAR (*17*). To determine the CTCF binding orientation (as shown in Fig. 2B) at CTCF peaks, FIMO (*18*) (threshold $P \leq 1$ e-4) was used to identify CTCF binding sites within the 50 bp flanking regions of ChIP summits of CTCF peaks that were co-occupied with top 500 ZFP661 peaks in ZFP661-3HA OE mESCs. The CTCF binding orientation was assigned when only one CTCF binding orientation was detected, or when the prediction score of one orientation was remarkably higher than that of the other orientation, in cases where both two CTCF binding orientations were present.

To determine the relative binding locations of CTCF, RAD21 and ZFP661 on CTCF-ZFP661 co-occupied peaks, the ChIP summits of each protein were used to indicate their binding positions. The relative binding positions of "ZFP661 to CTCF" and "RAD21 to CTCF" in each biological replicate were calculated. Only the peaks with the identical side for location relative to CTCF in the two biological replicates were retained and the average were taken for relative locations. In addition, the binding sites predicted using binding motifs were also used to determine the location of ZFP661 binding relative to CTCF binding. CTCF binding sites and ZFP661 binding sites were identified within the 50 bp franking regions of summits of CTCF peaks that were co-occupied with ZFP661 peaks using FIMO (threshold $P \leq 1$e-4) with CTCF binding motif (MA0139.1) and ZFP661 binding motif derived from ChIP-seq data, respectively. When two or more binding sites were identified for CTCF or ZFP661, the binding site with the highest score was used. Then, the relative positions of ZFP661 binding sites to CTCF binding sites were determined.

Hi-C and data analyses

Hi-C assays were performed using ZFP661-3HA OE mESCs. Cells were crosslinked with 2% formaldehyde for 10 min and quenched using 0.4M glycine for 5 min. Cells with approximately 4 µg of DNA were used as input to generate proximally ligated DNA using the Arima-HiC kit (Arima, A510008) according to the manufacturer's protocol (Arima, A160134 v01). Briefly, crosslinked cells were digested using the Arima restriction enzyme cocktail, the 5' overhangs were filled in and labeled with a biotinylated nucleotide, and the spatially proximal ends were ligated. The proximally ligated DNAs were purified and sonicated using the Covaris ME220 (8 microTube-130 AFA fiber strip V2: duration 42s, peak power 70w, duty factor 20%, cycle per burst 1000#). Then, size selection was performed on the resulting DNA Fragments (for size ranging from 200 to 600 bp). 2 µg of the resulting DNA fragments were used as input, and fragments labelled with biotin were bound to beads to construct Hi-C libraries using the KAPA HyperPrep Kit (Roche, KK8502) following the protocol provided by Arima (A160139 v00). The resulting libraries were sequenced using NovaSeq 6000 (paired-end 2x100 bp).

For data analysis, mouse genome (mm10) fragments were generated using the "digest_genome" tool in HiC-Pro utilities (*19*) with restriction motifs "GATC" and "G^ANTC". Raw fastq reads were cleaned using Trimmomatic. HiC-Pro pipelines (v2.11.4) (*19*) were then used with above genome fragments and the parameter "LIGATION_SITE = GATCGATC, GANTGATC,

GANTANTC, GATCANTC” to generate valid interaction pairs and contact maps. Briefly, the clean reads were mapped onto the mouse genome (mm10), and all valid interaction pairs were parsed with duplicates removed. Valid pairs from the two biological replicates in the same condition (*i.e.*, EV control or ZFP661-3HA OE) were pooled together, and pooled valid interaction pairs were used to generate corresponding ICE normalized Hi-C contact matrices.

GENOVA (*20*) was used to calculate insulation scores at CTCF barriers with the parameter “window = 5”, and normalized contact matrices with 1 kb resolution for EV control and ZFP661-3HA OE were used as input, respectively. CTCF binding sites that were not co-occupied with ZFP661 were used as controls. Single-tailed paired t-tests were performed to determine whether ZFP661 binding decreased the insulation function (correlating with higher insulation score) of CTCF barriers.

Chromatin loops were detected from both 5 kb-resolution and 10 kb-resolution Hi-C maps using HiCCUPS in Juicer (v1.6) (*21*) with the following parameters: “-r 5000,10000 -k KR -f .1,.1 -p 4,2 -i 7,5 -t 0.02,1.5,1.75,2 -d 20000,20000 --ignore-sparsity”, and pooled valid interaction pairs in EV control were used as input. The loop anchors from 5 kb maps (where the length of anchors equaled 5 kb) were extended to 10 kb by extending 2.5 kb on each side. Loops with one anchor (referred to as “Anchor B”) overlapping with CTCF barriers (meeting the following requirements simultaneously: the orientation of CTCF binding site was towards inside the loops and ZFP661 binding was inside the CTCF barriers) were extracted for downstream analyses. To minimize Hi-C background noise, only loops with the other loop anchor (referred to as “Anchor A”) located no less than 100 kb from “Anchor B” were retained. Valid interaction pairs with one end located at “Anchor A” and the other end located within the 50kb flanking regions of “Anchor B” were extracted in EV control and ZFP661-3HA OE samples, respectively. Then, a bin size of 5 kb with a sliding step of 2.5 kb was used to count the number of resulting interaction pairs with one end located within each bin for EV control and ZFP661-3HA OE samples, respectively. A scale number was calculated to make the number of interaction pairs with one end located in the 50 kb flanking regions of CTCF barriers equal between EV control and ZFP661-3HA OE, and the number of interaction pairs in each bin was then normalized by multiplying it with this scale number. Finally, a $\log_2$(ZFP661 OE/EV) was calculated in each bin surrounding CTCF barriers that were co-occupied with ZFP661 to check whether the chromatin loop was trapped at CTCF barriers or passed through them. The curve displayed was smoothed using the spline function in R.

<u>GO enrichment analysis</u>
The gene annotation for mouse genome (mm10) was obtained from GENCODE (*22*) (M23, only protein coding genes on the reference chromosomes were used). ZFP661 target genes (n = 473) were assigned based on their proximity to ZFP661 binding peaks (top 500 in ZFP661-3HA OE mESCs): genes with at least one ZFP661 peak located within the 10 kb flanking regions of their gene bodies (from TSS to TES) or at their gene bodies were considered as ZFP661 target genes. GO enrichment analysis was performed using DAVID (v2021) (*23*) on “biological processes” terms (annotation v2022q3; threshold: FDR-adjusted $P \leq 0.1$).

<u>Capture Hi-C and data analyses</u>

Timed mouse breeding (*Zfp661*$^{+/-}$ x *Zfp661*$^{+/-}$) was set up in the early evening, and plugs were checked in the following early mornings. The morning when the plug was observed was marked as E0.5, and female mice with plugs were removed and housed in new cages. Embryonic forebrains were isolated at E16.5 and placed on ice during genotyping. Genotypes were quickly determined using the Platinum Direct PCR Universal Master Mix (Invitrogen, A44647100). Embryonic forebrains from *Zfp661*$^{+/+}$ and *Zfp661*$^{-/-}$ littermate pairs were dissociated into a single-cell suspension using the Papain Dissociation System (Worthington LK003150) by incubating 45 min at 37°C. The cells from E16.5 forebrains were then processed using the same protocol as mESCs (described in the "Hi-C and data analyses" section) to construct Hi-C libraries. Three biological replicates for each genotype were used.

To capture the three Pcdh clusters and nearby genomic regions (mm10, chr18: 36,630,000-38,080,000), probes were designed as follows: for each cutting site on the target region, a buffer of 70 bp was assigned on each side, and the next 60 bp in each direction were designated as probe seed regions (with a probe length of 120 bp); all probe seed regions within the capture region were then combined and used as input for the SureSelect DNA tool of Agilent SureDesign to generate a probe set, with the following settings: 2X tiling density, moderately stringent masking, and optimized performance for HS2/XT. The resulting probe set was used for Capture Hi-C assays.

Capture assays were performed using the above probe set with the Hi-C libraries of E16.5 forebrains and ZFP661-3HA OE mESCs as inputs, respectively. The Capture Hi-C libraries were obtained by fast hybridization for 90 min with probes, followed by capture, washes and post-capture amplification using the SureSelect XT HS2 DNA Target Enrichment Kit (Agilent, G9987A) according to the protocol (Agilent, G9983-90000). The resulting Capture Hi-C libraries were sequenced using NovaSeq 6000 (paired-end 2x100bp). The raw reads were cleaned using Trimmomatic, and the clean reads were further processed using HiC-Pro (v2.11.4) to obtain valid interaction pairs following the protocol described in the "Hi-C and data analyses" section. Valid interaction pairs from the biological replicates of the same condition (*i.e.*, *Zfp661*$^{+/+}$ or *Zfp661*$^{-/-}$ E16.5 forebrains, EV control or ZFP661-3HA OE) were combined, and only the interaction pairs where both ends were located within the capture regions (chr18: 36,630,000-38,080,000) were retained for downstream analyses. The resulting interaction pairs were used to generate raw contact matrices using HiC-Pro.

The position information of the enhancers downstream of the *Pcdhγ* locus was obtained from the "ENCODE Regulation Tracks" (*24*) on UCSC genome browser (*25*) (mm10, strong enhancers in E16.5 forebrains), which contained five strong enhancers with the coordinate spanning chr18: 37,839,600 - 37,880,000. Gene annotation for Pcdhs was obtained from GENCODE (M23), and only protein-coding genes were retained for downstream analyses. Using 5 kb-resolution raw contact matrices as input, GENOVA was used to visualize the Hi-C map in *Zfp661*$^{+/+}$ E16.5 mouse forebrains, and the differential Hi-C maps between *Zfp661*$^{-/-}$ and *Zfp661*$^{+/+}$ in E16.5 forebrains, and between ZFP661-3HA OE and EV controls in mESCs. The region displayed in the differential Hi-C maps covered the *Pcdhβ and Pcdhγ* loci and the downstream enhancer cluster (mm10, chr18: 37,260,000-37,885,000). To reduce possible noise, interaction pairs with a distance less than 20 kb were filtered out, and the samples were scaled to the same number of interaction pairs during GENOVA loading the contact matrices.

To determine the contact between downstream enhancers and each *Pcdhβ/γ* promoter (from Pcdhb1 to Pcdhga12), the interaction pairs with one end located at the enhancers and the other end located at the promoter regions (TSS ± 2.5kb) were extracted and counted. The count numbers were further normalized by scaling the total contact numbers in the sample with lower counts to that of the sample with higher count between *Zfp661*$^{+/+}$ and *Zfp661*$^{-/-}$ in E16.5 forebrains, and between EV control and ZFP661-3HA OE in mESCs, respectively. The isoforms with total interaction number less than 100 interaction pairs (less than 0.8% of total enhancer-promoter interaction pairs) in the sum of the two conditions (*Zfp661*$^{+/+}$ and *Zfp661*$^{-/-}$ in E16.5 forebrains, or EV control and ZFP661-3HA OE in mESCs) were not displayed.

<u>Single-cell 5' RNA-seq and assignment of cell types</u>
Timed mouse breeding (*Zfp661*$^{+/-}$ x *Zfp661*$^{+/-}$) was set up as described in the "Capture Hi-C and data analyses" section. At E16.5, embryonic forebrains were isolated and dissociated into single-cell suspension using the Papain Dissociation System by incubating at 37°C for 45 min. At the same time, the genotypes of the embryos were quickly determined using the Platinum Direct PCR Universal Master Mix. After the incubation, the cell suspension of two littermate pairs of *Zfp661*$^{+/+}$ and *Zfp661*$^{-/-}$ mice was passed through cellTrics Cell Strainers (20 µm) to remove cell debris and clumps. The suspension was then subjected to a discontinuous density gradient centrifugation using the Papain Dissociation System at 70 g for 6 min to further remove remaining debris and membrane fragments. The cells were collected, resuspended in 0.04% BSA/PBS solution, and washed twice using 0.04% BSA/PBS solution.

The resulting single-cell samples were counted using the Countess II Automated Cell Counter, and the cell concentration was adjusted to 1000 cells/µl for single-cell barcoding. The samples were barcoded using the 10x Chromium Controller by targeting for 2000 recovered cells per sample to decrease the multiplet rate. Single-cell 5' RNA-seq libraries were constructed using Chromium Next GEM Single Cell 5' Reagent Kits v2 (Dual Index) (10x, PN-1000265) following the manufacturer's protocol (10x, CG000331 Rev D). The resulting libraries were sequenced twice using NovaSeq 6000 (sequencing configuration: read 1: 28 bp; i7 index: 10 bp; i5 index: 10 bp; read 2: 150 bp) to ensure that the sequencing depth per cell amongst samples matched.

The gene annotation was obtained from GENCODE (M23), and only protein-coding genes were retained for downstream analyses. Raw fastq reads were mapped onto the mouse genome (mm10), and gene-cell matrices were generated using the count function in Cell Ranger (v7.0.0). High-quality cells were selected based on the following criteria: cells with 2000-40000 Unique Molecular Identifiers (UMIs) and 1000-6500 genes detected, and with the percentage of UMIs from mitochondrial genes no greater than 10%. The gene-cell matrices were updated by selecting high-quality cells and excluding mitochondrial genes using the reanalysis function in Cell Ranger. To further ensure that the distribution of UMIs/cell between *Zfp661*$^{+/+}$ and *Zfp661*$^{-/-}$ littermates were well matched, UMI counts within a cell were downsampled as follows: for each sample, cells were ranked based on the number of UMIs detected and divided into 20 equal parts. The downsampleMatrix function in R package "scuttle" (*26*) was then used to downsample UMI counts to ensure that the mean UMIs/cell in each part equaled the corresponding part in the matched littermate sample with a lower mean UMIs/cell. The resulting four samples (two littermate pairs) were integrated using Seurat (v4.3.0) (*27*) with the "vst" method selecting for top 2000 variable features.

To build a reference map for E16.5 mouse forebrains, raw sequencing data from 11 mouse forebrain samples at the age of E16-17 were downloaded from Sequence Read Archive (PRJNA637987). The data were from a study (*28*) that reported the single-cell atlas of developing mouse brains, and cell types were well annotated in the dataset. The raw sequencing data were mapped onto the mouse genome (mm10), and gene-cell matrices were generated using Cell Ranger. The gene-cell matrices were further updated by selecting well-annotated cells, and the resulting matrices of the 11 samples were integrated using Seurat with the same settings as processing our own samples. Cell types for our own samples were assigned by projection onto the reference map for E16.5 mouse forebrains using Seurat with the first 30 dimensions of PCA selected. UMAP projection was performed using the top 2000 variable features that were selected by the "vst" method and the first 20 dimensions of PCA. Cell types with less than 150 cells in the integrated sample (including 4 sub-samples) were assigned as "Others", which were not displayed in the UMAP visualization.

<u>The usage of Pcdh isoforms and calculation of Pcdh-based similarity scores and repulsive signals amongst neurons</u>

The analyses in this study focused on the two largest cell types in the population, namely "Cortical or hippocampal glutamatergic neurons" (referred to as "glutamatergic neurons") and "Forebrain GABAergic neuron" (referred to as "GABAergic neurons"). The usage of Pcdhβ/γ isoforms (from Pcdhb1 to Pcdhga12) were calculated within the glutamatergic neurons, GABAergic neurons and pooled neurons of these two types, respectively. The usage of an isoform represents the proportion of UMIs from this isoform in all the UMIs from Pcdhβ/γ isoforms (from Pcdhb1 to Pcdhga12). Pcdhb1 was not detected and therefore not displayed in the usage visualization.

The Pcdh-based similarity score amongst neurons represents the probability of a Pcdh isoform taken from one neuron matching an isoform taken from another neuron. To account for the sparseness of single-cell data, the following steps were employed to ensure the calculation of similarity scores were more robust: 1) in each neuron type (glutamatergic and GABAergic neurons), only cells with a number of UMIs from Pcdhβ/γ isoforms no less than the mean Pcdhβ/γ UMIs/cell of that cell type were retained for downstream analyses; 2) for each biological sample (two pairs of littermates, four samples), the usage of Pcdh isoforms (including Pcdhα, Pcdhβ and Pcdhγ) in each neuron type was calculated and treated as one pseudocount (*i.e.*, background), and was added to the UMI counts of corresponding Pcdh isoforms. Similarity scores between any pairs of neurons within glutamatergic neurons, GABAergic neurons, and between the two types were then calculated in each biological sample according to the definition, respectively.

To compare the difference in the similarity score between *Zfp661*$^{+/+}$ and *Zfp661*$^{-/-}$ neurons, subsampling was performed as follows: for similarity scores in each type (glutamatergic-glutamatergic, GABAergic-GABAergic, and glutamatergic- GABAergic) of each biological sample, 50% of the datapoints were randomly sampled without replacement, and a median similarity score of this subsample was taken. Then, the median similarity scores from two biological replicates were averaged. This subsampling process was repeated 1000 times, and the data displayed in the Fig. 4D represents the distribution of median similarity scores of these 1000 subsamples. The $P$ value was calculated based on these 1000 subsamples, and "$P < 0.001$" represents that the median similarity score amongst *Zfp661*$^{+/+}$ neurons was less than that of *Zfp661*$^{-/-}$ neurons in all 1000 subsamples.

Repulsive signals amongst neurons (or neurites) were defined as the probability of sequential matching of Pcdh isoforms during continuous comparison by stimulating the chain formation process of Pcdh molecules. Mathematically, the repulsive signal equals $S^{n}$, where $S$ represents the similarity score, and n represents the number of continuous comparison (*i.e.*, Pcdh tiling number). For each subsample, the ratio of median similarity score between *Zfp661*$^{+/+}$ and *Zfp661*$^{-/-}$ neurons was calculated as $S_{WT}/S_{KO}$, and the ratio of repulsive signals under different Pcdh tiling number would be $(S_{WT}/S_{KO})^{n}$. The ratio of repulsive signals under different Pcdh tiling number (n = 1 to 10 shown in Fig. 4E) for 1000 subsamples were calculated, and the data were shown as mean ± SD in Fig. 4E.

Golgi-Cox staining, imaging and 3D reconstruction of neuronal dendrites

Three pairs of *Zfp661*$^{+/+}$ and *Zfp661*$^{-/-}$ littermates, including both male and female pairs aged between 60 to 66 days, were used in this assay. The brains were isolated and processed for Golgi-Cox staining using FD Rapid GolgiStain Kit (FD NeuroTechnologies, PK401) according to the manufacturer's protocol. During this process, brains were immersed in Solution A/B for 14 days, followed by Solution C for 7 days. Forebrains were sectioned using a Leica Cm3050s cryostat with a thickness of 100 µm. Serial coronal sections were collected, airdried and stained.

The experimenter who performed neuronal imaging, tracing and measurement was blinded to the genotype of mice. The brain sections in the Bregma 1.70 mm to 1.18 mm regions were selected, and the sections between littermate pairs were well-aligned based on the structures of the corpus callosum and striatum. To ensure consistency in neuron type and location during neuron sampling, pyramidal neurons in the somatomotor area of Layer 2/3 within approximately 300 µm from the brain surface (about half the radium of the field of view under 20X microscope) were selected. Neurons whose dendrites could be clearly traced were selected for imaging, and neurons were sequentially sampled from the middle to the sides of brain sections, with 8 neurons sampled per mice. Z-stack images were acquired to cover all visible dendrites using a Leica DM6000B (20X/0.70) microscope and LAS X software with optimal z-step of 0.52 µm and the same settings for all imaging.

To better trace neuronal dendrites from Golgi-Cox staining, Z-stacked images were pre-processed using Fiji/ImageJ2 (v2.9.0/1.53t) (*29*) as follows: the image format was transformed into 8-bit type, both the signals and LUT were inverted, and the scale unit was set to microns. All dendrites from sampled neurons were traced using the SNT toolbox (*30*) in Fiji/ImageJ2 with the "A* search algorithm" enabled, and the trace for each dendrite was further manually verified. Using the resulting 3D neuronal traces as input, Sholl analysis was performed using the SNT toolbox with a 10 µm radius step and within 150 µm from the soma. The branch number and total branch length of sampled neurons were also measured using the SNT toolbox. One-way repeated measures ANOVA was performed to determine whether there was a difference in the dendritic distributions of neurons from *Zfp661*$^{+/+}$ and *Zfp661*$^{-/-}$ mice. Single-tailed Wilcoxon rank-sum tests were performed to determine whether there were less branch number and total branch length in *Zfp661*$^{-/-}$ neurons than in *Zfp661*$^{+/+}$ neurons, respectively.

Sociability tests

Six pairs of *Zfp661*$^{+/+}$ and *Zfp661*$^{-/-}$ littermate mice, consisting of three pairs of males and three pairs of females aged between 2 to 4 months, were sent to the NHLBI Murine Phenotyping Core for sociability tests. The sociability tests were conducted in a rectangular three-compartment chamber with removable doors (Ugo Basile, 60 x 40 x 25 cm (h) Mouse Sociability Apparatus). The test mouse was allowed to explore the empty chamber freely for 6 min for habituation, after which it was placed in the central compartment with the removable doors closed. A stranger mouse of the same sex, which had been acclimated to being placed inside the grid enclosure previously, was placed inside the grid enclosure in one of the outer side compartments, while the other outer side compartment contained an empty grid enclosure. The test mouse was then permitted to explore the chamber freely for 6 min by lifting the removable doors. The tests were recorded, and the instances of the test mouse directly sniffing the stranger mouse were manually counted. The stranger mouse position was rotated in the outer side compartment (left or right) between tests.

After receiving the results of sociability tests from the Murine Phenotyping Core, a single-tailed paired t-test was performed to determine whether *Zfp661*$^{-/-}$ mice showed fewer directly sniffing interactions compared with *Zfp661*$^{+/+}$ mice. To improve visualization, one of the two data points that completely overlapped was slightly shifted by 0.5 sniffing interactions.

Conservation analysis of ZNF2/ZFP661 bindings in humans and mice

ZNF2 and ZFP661 are orthologs in humans and mice, respectively. Using the same approach as for identifying ZFP661 binding motif, MEME-ChIP was used to identify ZNF2 binding motif within the 50 bp flanking regions of the ChIP summits of the top 200 ZNF2 binding peaks that did not overlap with any CTCF peaks from ENCODE (v2). Tomtom was used to compare the ZNF2 and ZFP661 binding motifs derived from ChIP-seq data with default parameters. Gene annotation for the human genome (hg19) was obtained from GENCODE (v19). The top 500 binding peaks of ZFP661 were projected onto the human genome (hg19) using LiftOver (*25*) with default parameters.

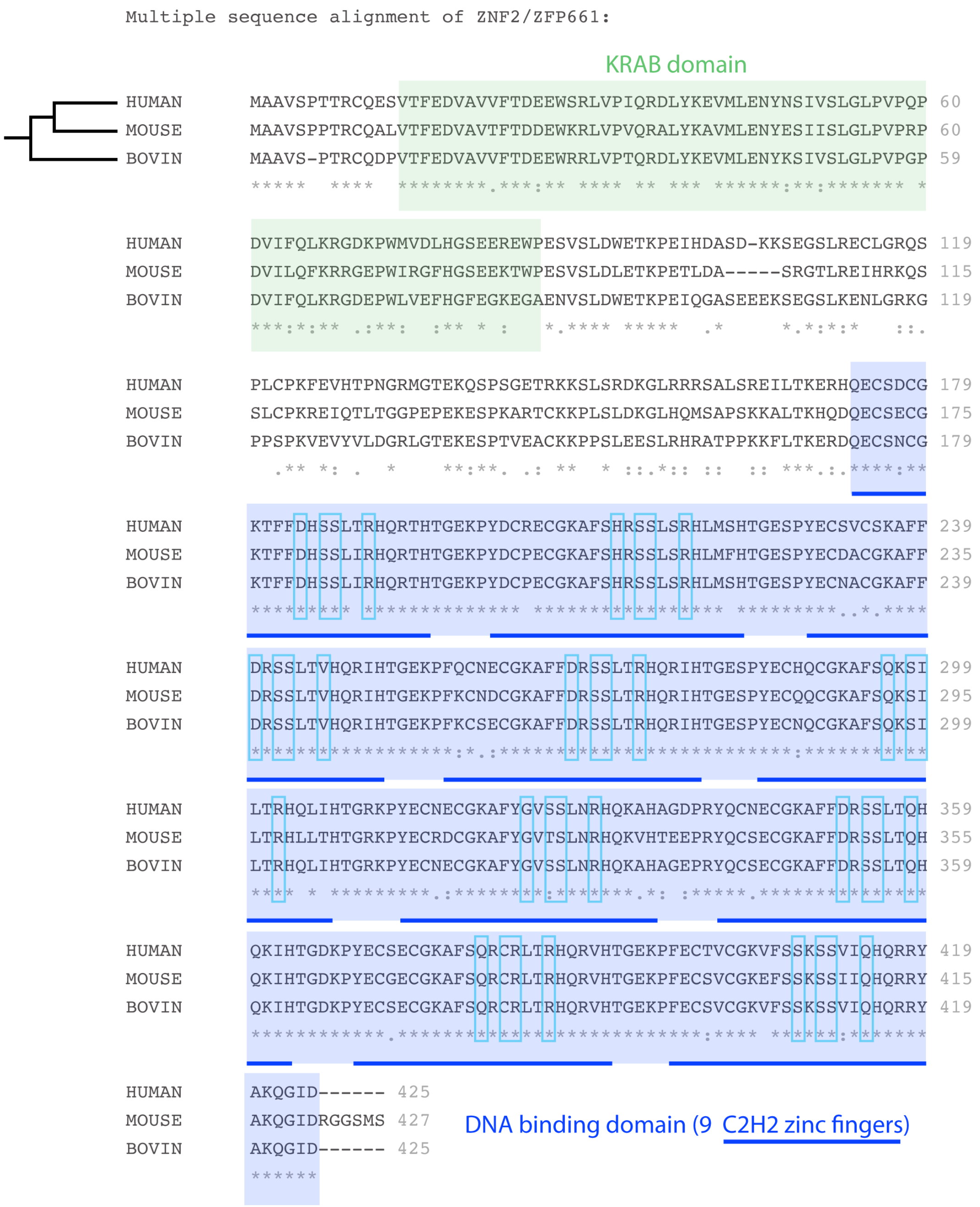


**Fig. S1. The DNA binding domain of ZFP661/ZNF2 is highly conserved.**
Multiple sequence alignment of Zfp661 orthologs was performed on three representative species: human (*Homo sapiens*), mouse (*Mus musculus*) and bovine (*Bos taurus*). The KRAB domain and DNA binding domain (DBD) are highlighted as indicated, respectively. The zinc fingerprint amino acids, which are responsible for making specific DNA base contacts, are framed by cyan rectangles. 90.1% (228/253) of the DBD sequences and 97.2% (35/36) of the fingerprint amino acids are identical across the three species.

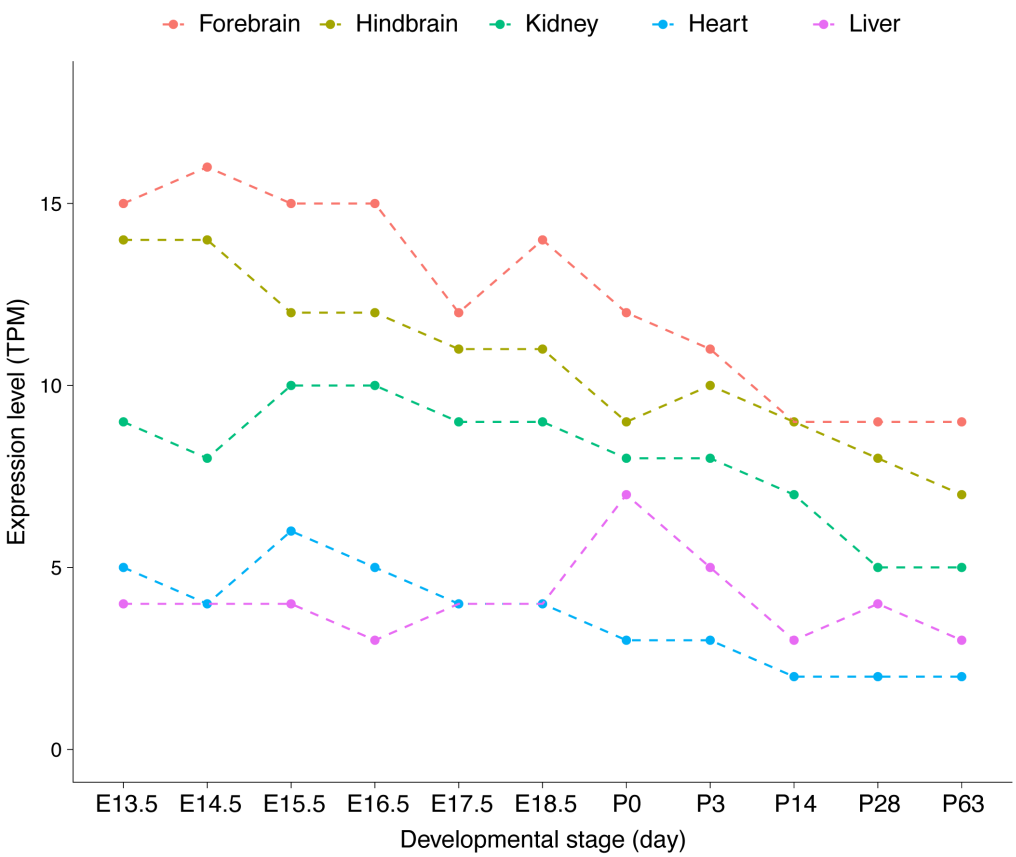


**Fig. S2. ZFP661 is relatively highly expressed in the developing brain of mice.**
The expression levels of Zfp661 mRNA in the forebrain, hindbrain, kidney, heart, and liver of mice at embryonic and postnatal stages are presented as transcripts per million (TPM). "E" and "P" represent embryonic day and postnatal day, respectively. The expression data were obtained from the Expression Atlas (*31*).

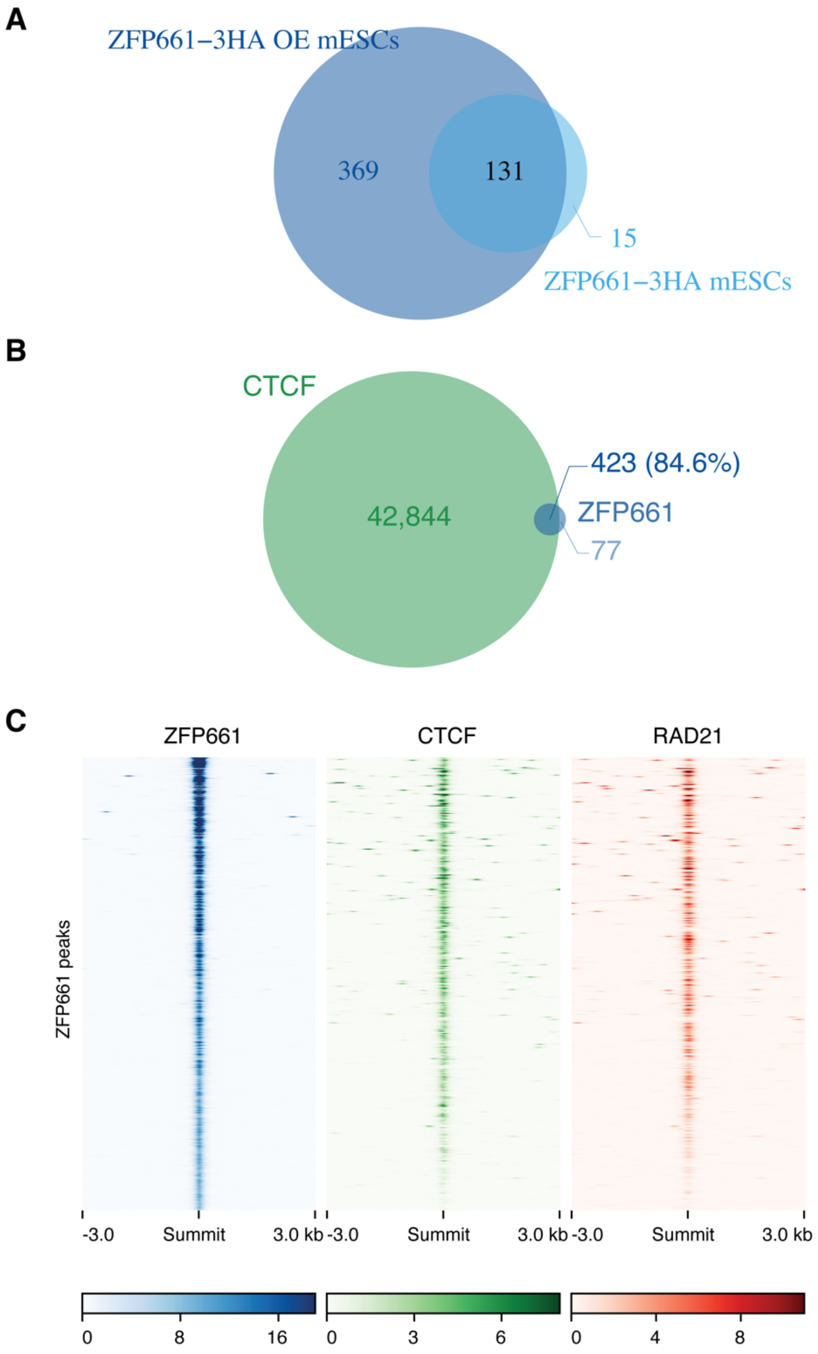


**Fig. S3. Overlap of ZFP661 binding peaks with CTCF and RAD21 in ZFP661-3HA OE mESCs.**
(**A**) Venn diagram shows that the top 500 ZFP661 binding peaks in ZFP661-3HA OE mESCs cover 89.7% (131 of 146) ZFP661 peaks detected in endogenous tagged ZFP661-3HA mESCs. (**B**) Venn diagram shows that 84.6% (423 of 500) of top 500 ZFP661 binding peaks overlap with CTCF binding peaks. (**C**) Heatmap shows the co-occupancy of ZFP661 peaks with those of CTCF and RAD21.

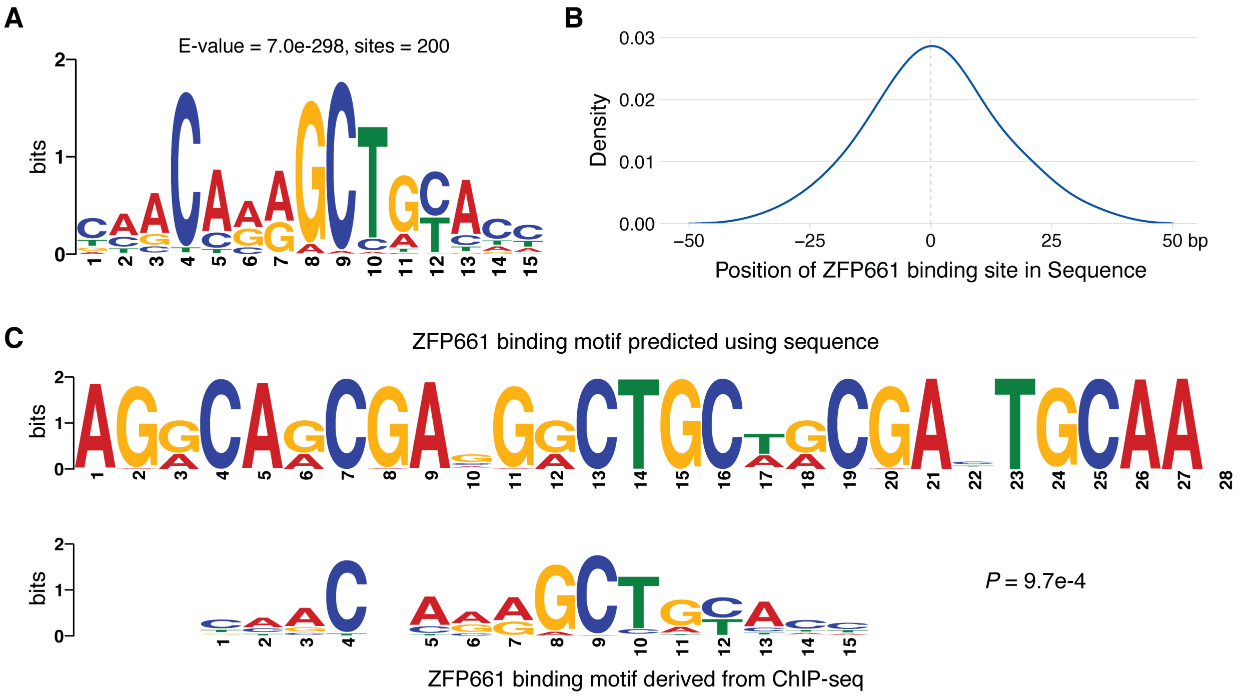


**Fig. S4. ZFP661 binding motif derived from ChIP-seq data closely matches the *de novo* predicted motif.**
(**A and B**) ZFP661 binding motif derived from the top 200 ZFP661 ChIP-seq peaks (summits ± 50bp) that do not overlap with CTCF peaks in ZFP661-3HA OE mESCs (A) and the distribution of this motif within the sequences (B). All 200 peak regions used to detect ZFP661 binding motif contain this motif. (**C**) Comparison between the ZFP661 binding motif derived from ChIP-seq data and the motif predicted *de novo* using the ZFP661 protein sequence (at website http://zf.princeton.edu/) (*14*).

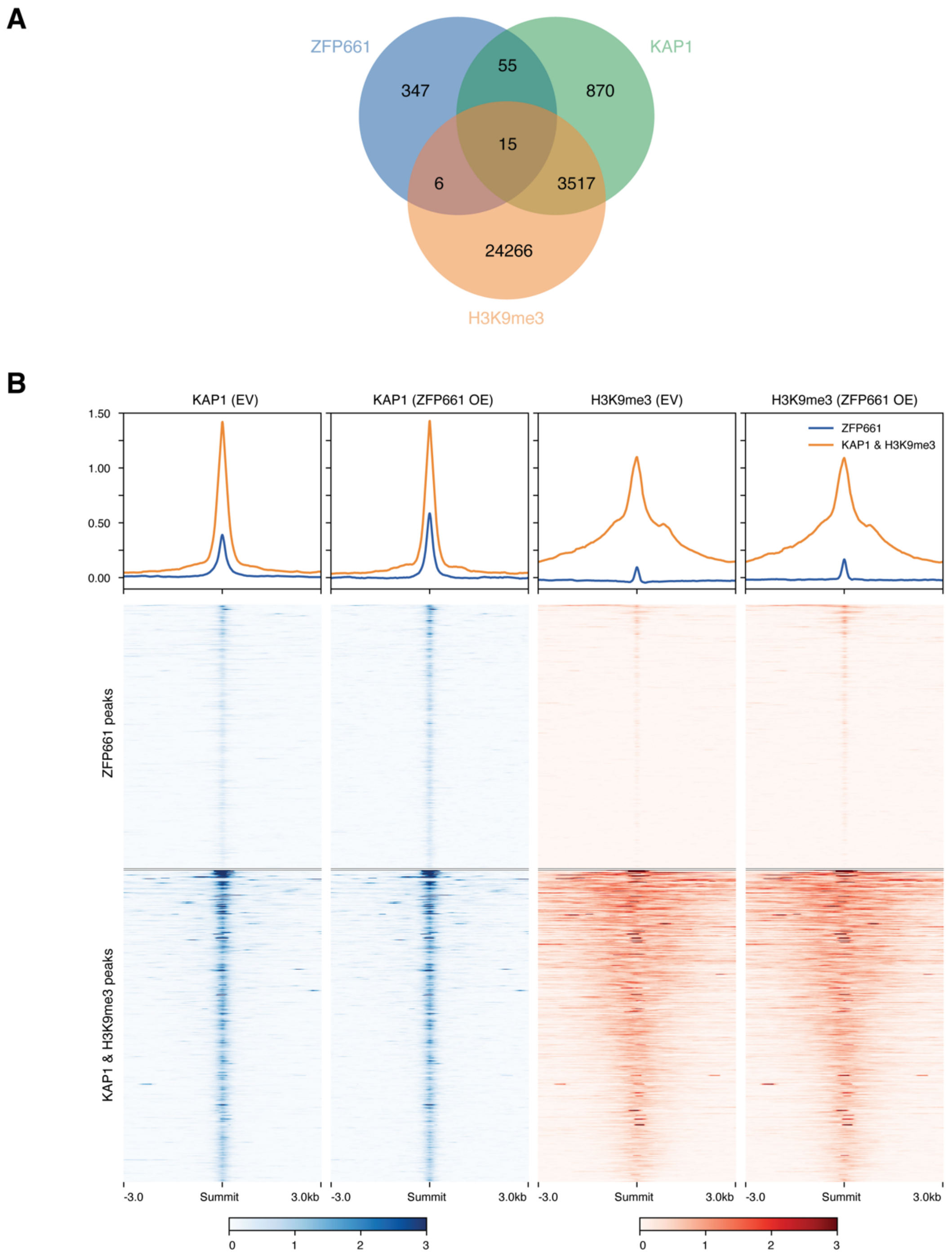


**Fig. S5. ZFP661 presents a weak capability to recruit KAP1 and establish H3K9me3.**
(**A**) Venn diagram shows the overlap amongst ZFP661, KAP1 and H3K9me3 peaks in ZFP661-3HA OE mESCs. Only 4.96% (21 of 423) of ZFP661 binding peaks are co-occupied with H3K9me3. (**B**) Alterations of KAP1 and H3K9me3 signals on ZFP661 peaks that are co-occupied with CTCF after overexpression of ZFP661. 500 randomly sampled peaks from the peaks co-occupied by KAP1 and H3K9me3 are used as controls.

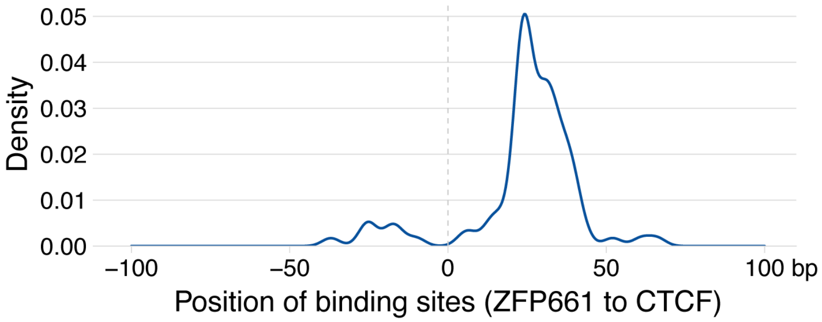


**Fig. S6. ZFP661 binding sites are located inside CTCF loop anchors.**
The relative locations of ZFP661 binding sites to CTCF binding sites are shown. ZFP661 and CTCF binding sites were identified by scanning ZFP661 and CTCF binding motifs on ZFP661-CTCF co-occupied peaks, respectively. CTCF binding orientation is as shown in Fig. 2B.

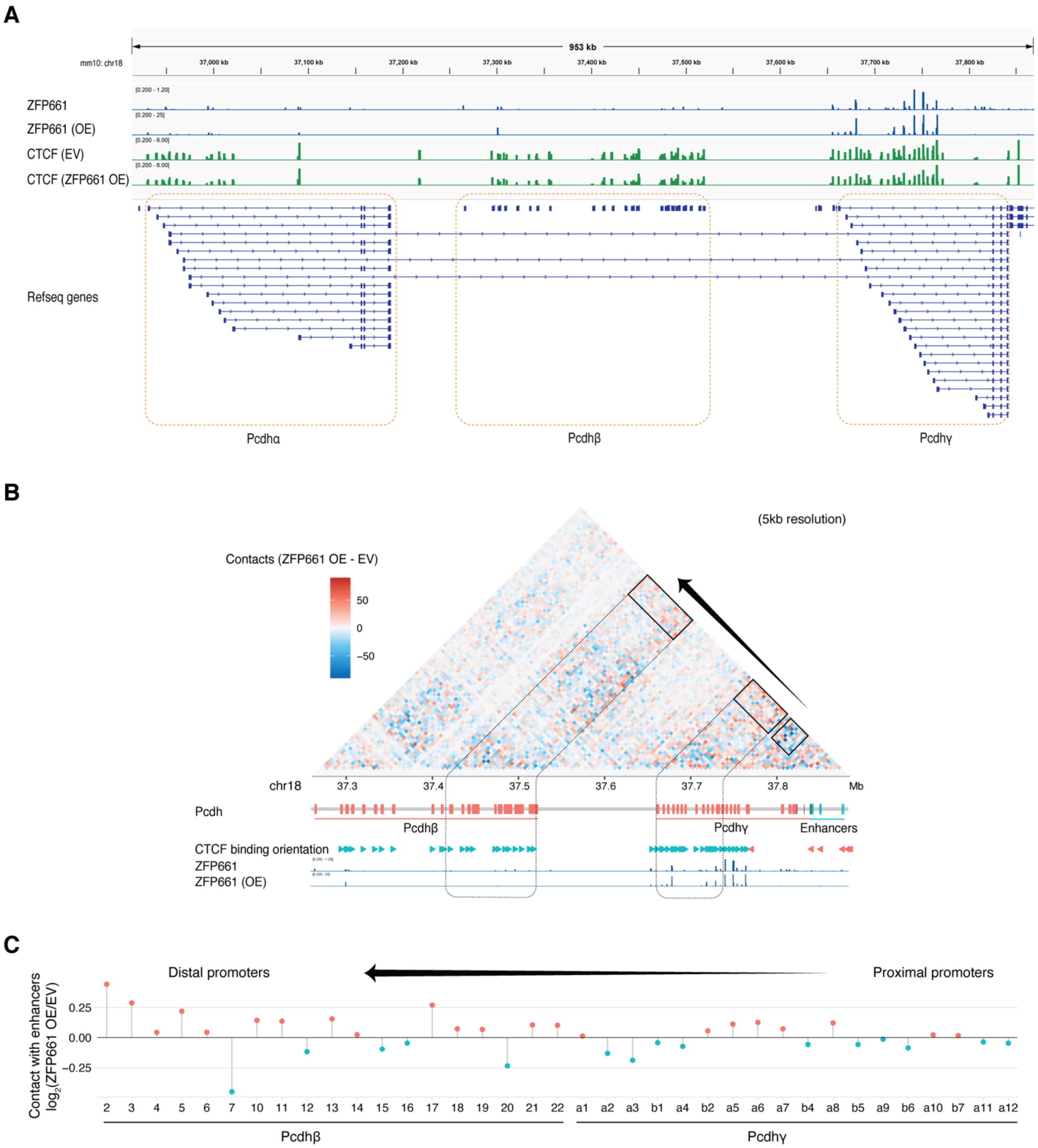


**Fig. S7. Overexpression of ZFP661 promotes the interaction of *Pcdhγ* downstream enhancers with distal *Pcdh* promoters.**

(**A**) Three protocadherin clusters in the mouse genome (mm10) with ZFP661 and CTCF occupancies shown in the upper portion. (**B and C**) Alterations of the contacts at the *Pcdhβ* and *Pcdhγ* regions (B) and the contacts between the enhancer cluster and individual promoters (TSS ± 2.5 kb) (C) after overexpression of ZFP661 in mESCs.

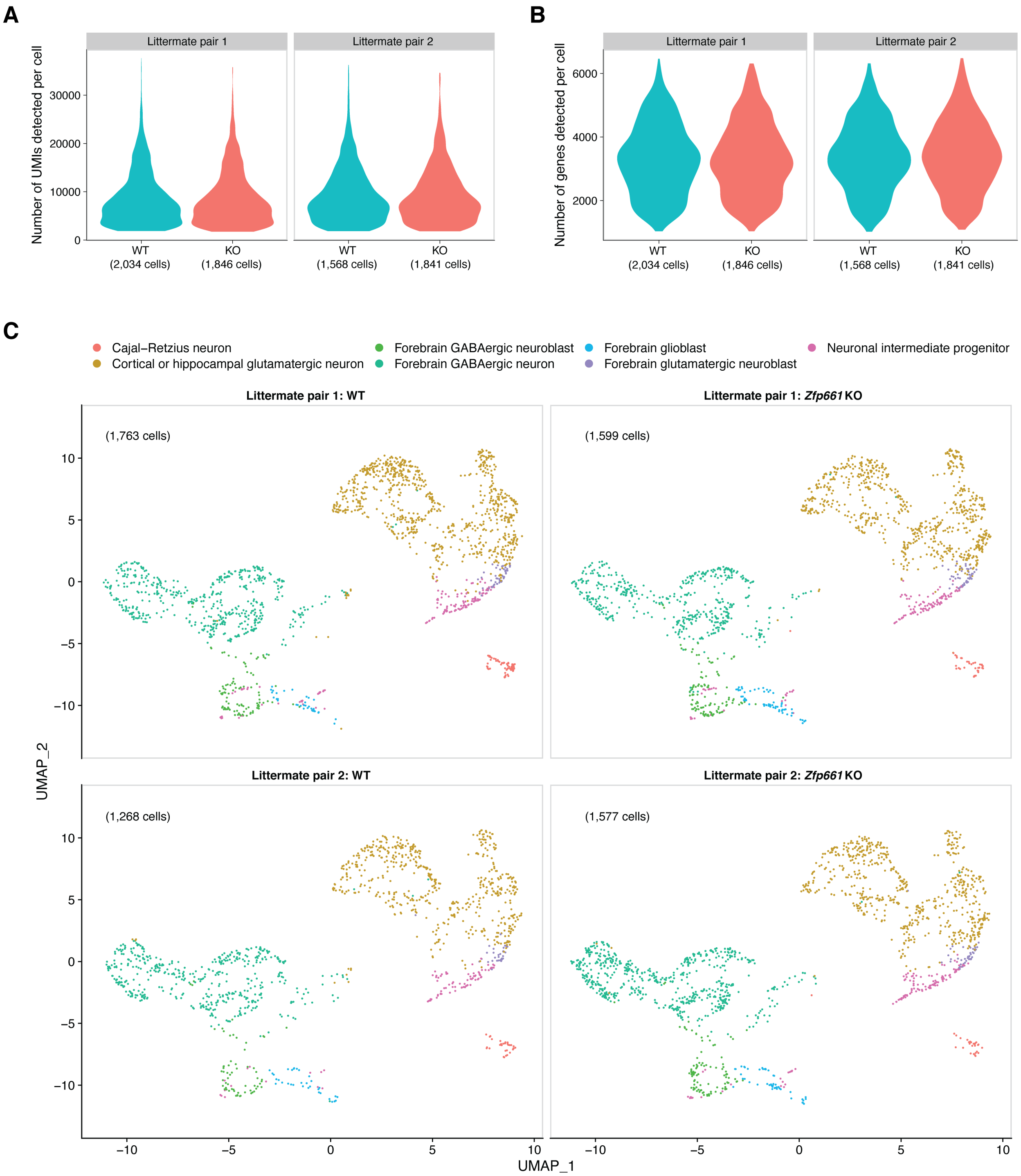


**Fig. S8. Single-cell atlas of *Zfp661*$^{+/+}$ and *Zfp661*$^{-/-}$ E16.5 mouse forebrains.**
(**A and B**) The distribution of unique molecular identifier (UMI) counts detected per cell (A) and the number of genes detected per cell (B). These features are well matched between *Zfp661*$^{+/+}$ and *Zfp661*$^{-/-}$ samples in each littermate pair. (**C**) Uniform Manifold Approximation and Projection (UMAP) visualization of four samples (two pairs of *Zfp661*$^{+/+}$ and *Zfp661*$^{-/-}$ littermates). Cell types were assigned by projection onto a reference brain map of the same ages (*28*).

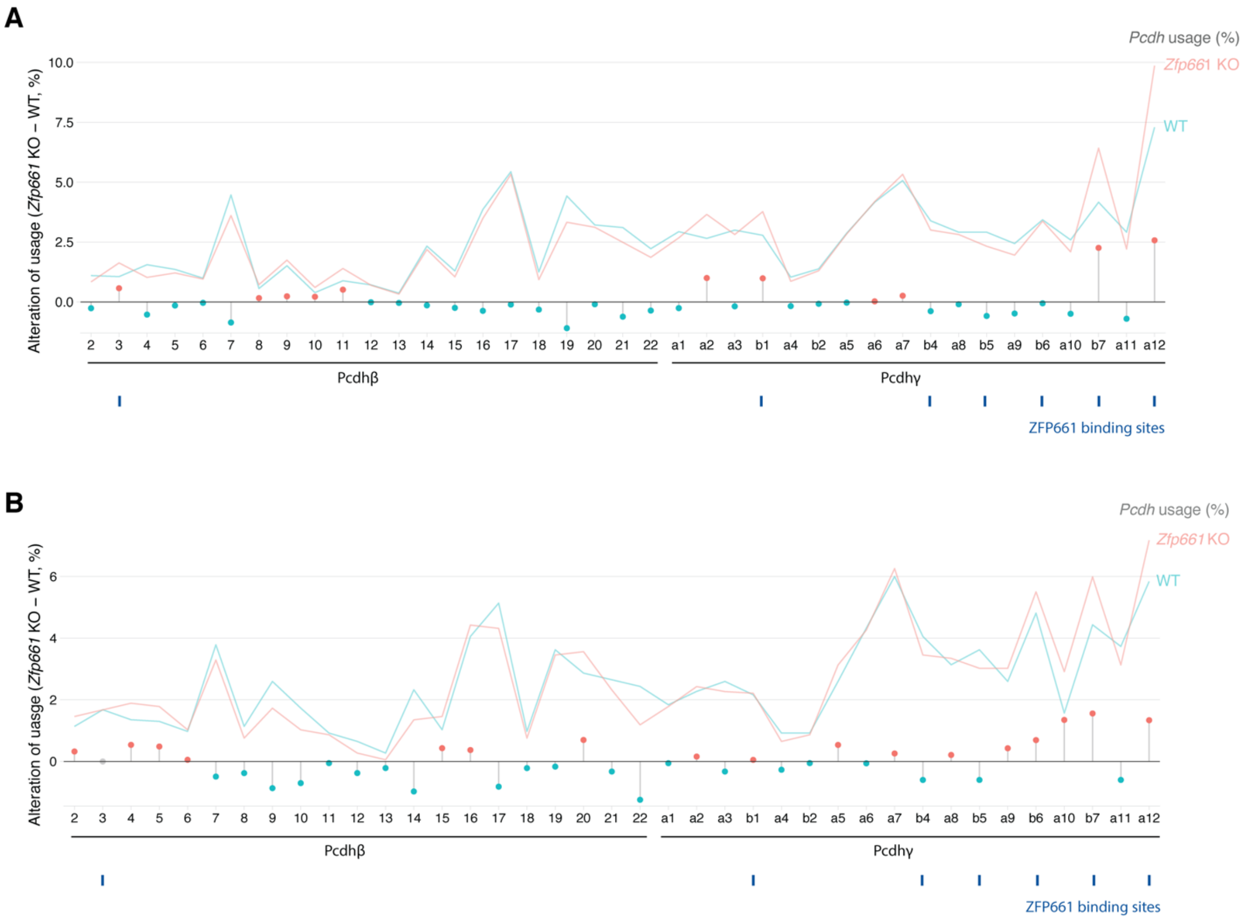


**Fig. S9. ZFP661 promotes usage of distal Pcdh isoforms in both glutamatergic and GABAergic neurons.**
(**A and B**) Alteration in usage of Pcdhβ/γ isoforms in glutamatergic neurons (A) and GABAergic neurons (B) after knocking out *Zfp661* in E16.5 mouse forebrains. The lines represent the level of usage and lollipops indicate the change in usage between *Zfp661*$^{+/+}$ and *Zfp661*$^{-/-}$ neurons.

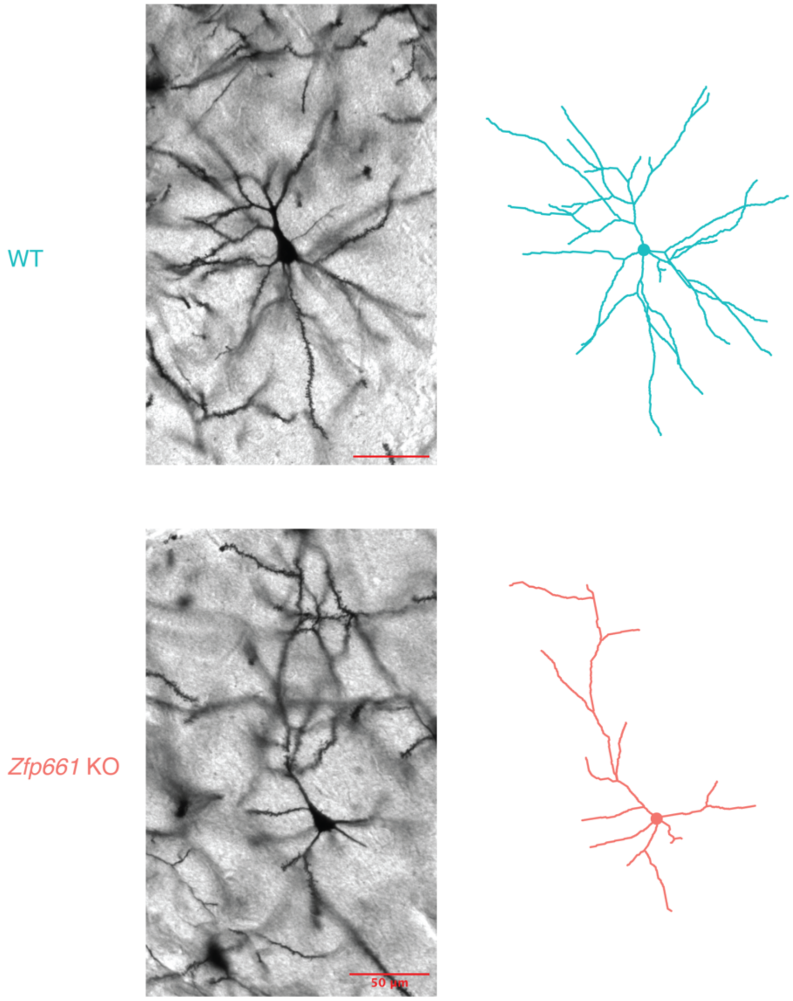


**Fig. S10. Representative microscopic images of Golgi-Cox stained neurons and 3D reconstructions of *Zfp661*$^{+/+}$ and *Zfp661*$^{-/-}$ neurons in mouse cerebral cortex.**
The neurons were sampled from layer II/III of the somatomotor areas of the cerebral cortex (Bregma: 1.70 to 1.18 mm) in early adult mice (P60-66). The two examples are at the same ranks based on the total branch length in the sampled *Zfp661*$^{+/+}$ and *Zfp661*$^{-/-}$ neurons.

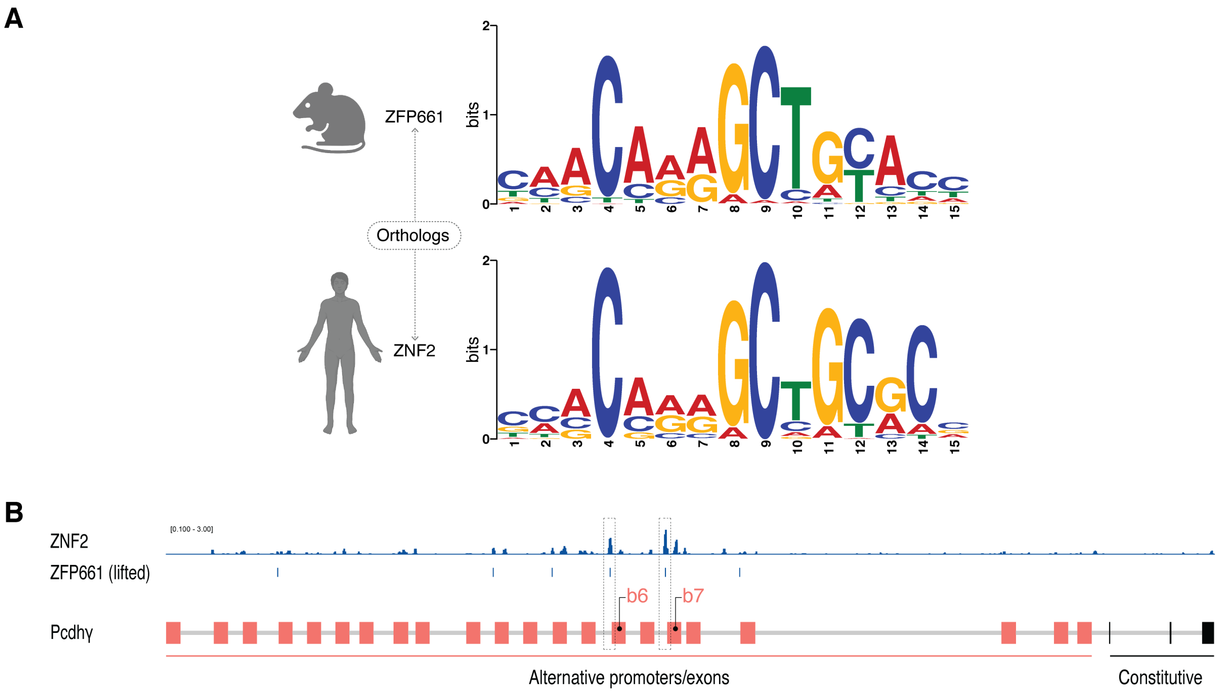


**Fig. S11. ZNF2/ZFP661 bindings are conserved between humans and mice.**
(**A**) Comparison of ZNF2 and ZFP661 binding motifs derived from ChIP-seq. The binding motifs are highly conserved between humans and mice. (**B**) The bindings of ZNF2 and ZFP661 on the *Pcdhγ* locus are conserved between humans and mice. The binding sites of ZFP661 were mapped from mouse genome (mm10) onto human genome (hg19) using LiftOver.

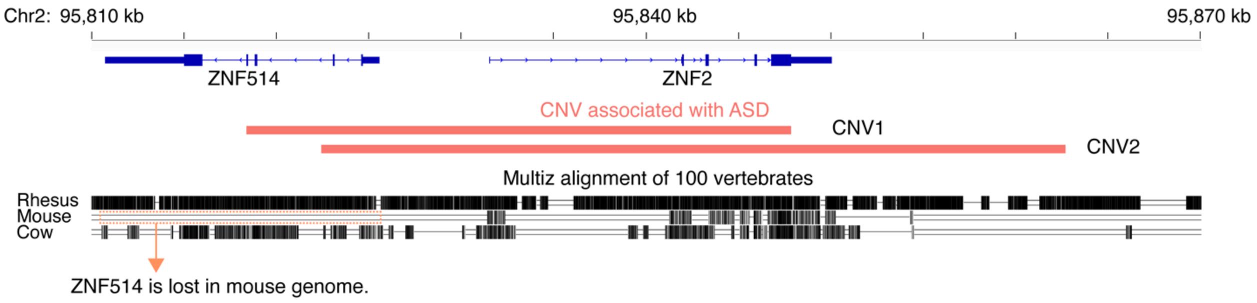


**Fig. S12. ZNF2 is associated with autism spectrum disorder (ASD) in humans.**
Copy number variations (CNVs) covering ZNF2 in human have been linked to ASD patients. CNV1 and CNV2 are from two distinct studies (*32, 33*).

| Cell and mouse model | Guide sequence |
| --- | --- |
| ***Zfp661* knockout** | g1: TGCACCGTTTACTCGGACCC<br>g2: CCGGCCCACCTGTTAGTGTT |
| **ZFP661-3HA** | g1: GGGAAGCATGAGCTAGACAG |

**Table S1.** Guide sequences used to generate *Zfp661* knockout and endogenously tagged ZFP661-3HA mESCs and mouse models.

| Cell/mouse Model | Sequence |
|---|---|
| **ZFP661-3HA (ssODN)** | ACCCTTCCAGGCTCGCAGGAAGGCCCGCTGTCTAAGCGTAATCTGGAA<br>CATCGTATGGGTAAGCGTAATCTGGAACATCGTATGGGTAAGCGTAAT<br>CTGGAACATCGTATGGGTAGCTCATGCTTCCCCCTCTGTCTATCCCTTG |
| **ZFP661-3HA OE** | ATGGCAGCAGTTAGTCCACCGACTCGCTGCCAAGCTTTGGTAACCTTT<br>GAAGACGTGGCTGTCACATTTACCGACGATGAGTGGAAGCGGCTCGT<br>GCCAGTGCAAAGGGCACTTTATAAGGCAGTGATGCTCGAGAATTATG<br>AAAGTATCATCTCACTGGGTCTGCCAGTGCCAAGACCAGACGTTATTC<br>TCCAGTTTAAGCGCAGGGGCGAGCCCTGGATTAGGGGCTTCCACGGGT<br>CAGAGGAAAAGACTTGGCCAGAATCAGTGTCACTTGACCTTGAAACA<br>AAGCCCGAGACCCTGGACGCAAGCCGGGGGACCCTCCGGGAAATTCA<br>CAGAAAACAGAGTTCACTGTGCCCAAAGAGGGAGATCCAGACACTGA<br>CCGGCGGTCCAGAGCCTGAGAAGGAATCACCTAAGGCCCGAACTTGC<br>AAGAAGCCCCTTTCACTCGACAAAGGCCTGCATCAGATGAGTGCTCCC<br>TCCAAGAAAGCCTTGACCAAGCATCAAGATCAGGAATGTTCTGAATG<br>CGGGAAGACCTTCTTCGATCACAGCAGCTTGATTAGACACCAGAGGA<br>CACACACCGGGGAGAAACCCTACGACTGCCCCGAGTGCGGGAAAGCC<br>TTCAGTCACAGAAGTTCCCTGAGTAGACACCTGATGTTCCACACCGGC<br>GAGAGTCCTTACGAATGCGATGCATGTGGCAAGGCATTTTTCGACCGC<br>TCTTCACTGACAGTGCACCAAAGGATCCACACGGGTGAAAAACCTTTC<br>AAGTGCAATGACTGTGGAAAAGCTTTCTTCGATAGATCAAGCTTGACC<br>AGACATCAGAGGATACATACCGGCGAATCACCATACGAGTGTCAGCA<br>GTGCGGAAAAGCCTTCAGCCAGAAATCTATCCTGACTCGCCACCTCCT<br>CACTCACACTGGACGCAAGCCATACGAATGCAGAGACTGCGGGAAAG<br>CCTTTTACGGTGTGACAAGCCTGAATCGGCACCAGAAAGTGCATACCG<br>AGGAACCGAGGTACCAATGTAGCGAGTGCGGGAAGGCTTTCTTTGAC<br>AGGAGCTCCCTCACCCAGCATCAAAAAATTCACACAGGGGACAAGCC<br>ATACGAGTGTGGGGAGTGTGGAAAGGCTTTCTCACAGAGATGTCGGTT<br>GACTAGACATCAGAGGGTGCATACTGGGGAAAAGCCATTTGAATGTT<br>CTGTTTGCGGTAAGGAGTTTTCTAGCAAGTCTTCCATCATCCAGCATC<br>AGAGAAGGTACGCCAAACAGGGTATCGACAGGGGTGGATCCATGTCT<br>TACCCCTACGACGTGCCCGACTACGCCGGCTATCCGTATGATGTCCCG<br>GACTATGCAGGATCCTATCCATACGACGTTCCAGATTACGCT |

**Table S2.** Single-stranded oligodeoxynucleotides (ssODNs) used to generate endogenously tagged ZFP661-3HA mESCs and mouse models, and the sequence of ZFP661-3HA used to generate ZFP661 OE mESCs.

| Model | Primer | Genotyping |
| --- | --- | --- |
| ***Zfp661* knockout** | F: GCGGGAGTCTATCACTGAGC<br>R1: CGGCTCACGAGACATCAGAG<br>R2: CTAGCGTCTCTCCCCTCTGT | WT: F + R1 (386bp)<br>KO: F + R2 (~290bp) |
| **ZFP661-3HA** | F: CACCGTTTACTCGGACCCTG<br>R: GACGTTACGCCAAACAAGGG | WT: 172 bp<br>KI: 253bp |

**Table S3.** Primers used to genotype *Zfp661* knockout and endogenously tagged ZFP661-3HA mESCs and mouse models.
